# A chromosome-level genome of black rockfish, *Sebastes schlegelii*, provides insights into the evolution of live birth

**DOI:** 10.1101/527036

**Authors:** Yan He, Yue Chang, Lisui Bao, Mengjun Yu, Rui Li, Jingjing Niu, Guangyi Fan, Weihao Song, Inge Seim, Yating Qin, Xuemei Li, Jinxiang Liu, Xiangfu Kong, Meiting Peng, Minmin Sun, Mengya Wang, Jiangbo Qu, Xuangang Wang, Xiaobing Liu, Xiaolong Wu, Xi Zhao, Xuliang Wang, Yaolei Zhang, Jiao Guo, Yang Liu, Kaiqiang Liu, Yilin Wang, He Zhang, Longqi Liu, Mingyue Wang, Haiyang Yu, Xubo Wang, Jie Cheng, Zhigang Wang, Xun Xu, Jian Wang, Huanming Yang, Simon Ming-Yuen Lee, Xin Liu, Quanqi Zhang, Jie Qi

## Abstract

Black rockfish (*Sebastes schlegelii*) is a teleost species where eggs are fertilized internally and retained in the maternal reproductive system, where they undergo development until live birth (termed viviparity). In the present study, we report a chromosome-level black rockfish genome assembly. High-throughput transcriptome analysis (RNA-seq and ATAC-seq), coupled with *in situ* hybridization (ISH) and immunofluorescence, identify several candidate genes for maternal preparation, sperm storage and release, and hatching. We propose that zona pellucida (ZP) genes retain sperm at the oocyte envelope, while genes in two distinct astacin metalloproteinase subfamilies serve to release sperm from the ZP and free the embryo from chorion at pre-hatching stage. Finally, we present a model of black rockfish reproduction, and propose that the rockfish ovarian wall has a similar function to uterus of mammals. Taken together, these genomic data reveal unprecedented insights into the evolution of an unusual teleost life history strategy, and provide a sound foundation for studying viviparity in non-mammalian vertebrates and an invaluable resource for rockfish ecology and evolution research.

## Introduction

Viviparity – the process of internal fertilization of an egg, development in a parental reproductive system (usually maternal), and live birth – has evolved independently in diverse vertebrate groups^1,2^. It is rare in teleosts, ray-finned fishes, where only 500 out of 30,000 species employ this life-history strategy while the remaining species are egg-laying (oviparous)^3^. Viviparity has been reported in five teleost orders (Lophiiformes, Beloniformes, Cyprinodontiformes, Scorpaeniformes, and Perciformes)^4,5^. Previous studies in fish focused on species of the Poeciliidae family within Cyprinodontiformes^6–8^. Recent genomic analyses on platyfish (*Xiphophorus maculatus*) and swordtail (*Xiphophorus hellerii*), viviparous species in the Poeciliidae family, revealed positive selection of protein-coding genes associated with reproductive features^2,9^. Exploring genetic mechanisms in orders other than Cyprinodontiformes promises to further improve our understanding of viviparity.

Rockfish (order Scorpaeniformes) include both viviparous and oviparous species. Black rockfish (*Sebastes schlegelii*; hereafter denoted ‘rockfish’) (Fig. 1a) has evolved viviparity. Previous reports on its reproduction process^10–12^ lay an extensive understanding of viviparity. Yet the associated genetic mechanism remains unexplored. We report here a first chromosome-level whole-genome assembly for rockfish and the dissection of rockfish reproduction – from mating to hatching – by integrating RNA-seq and ATAC-seq data, *in situ* hybridization (ISH), and immunofluorescence. From this dataset, we were able to identify crucial genes and gene families related to viviparity, especially in the stages sperm storage, pre-fertilization and hatching – providing an unprecedented genome-wide view of an unusual reproductive mode in teleost fishes.

**Fig. 1.**
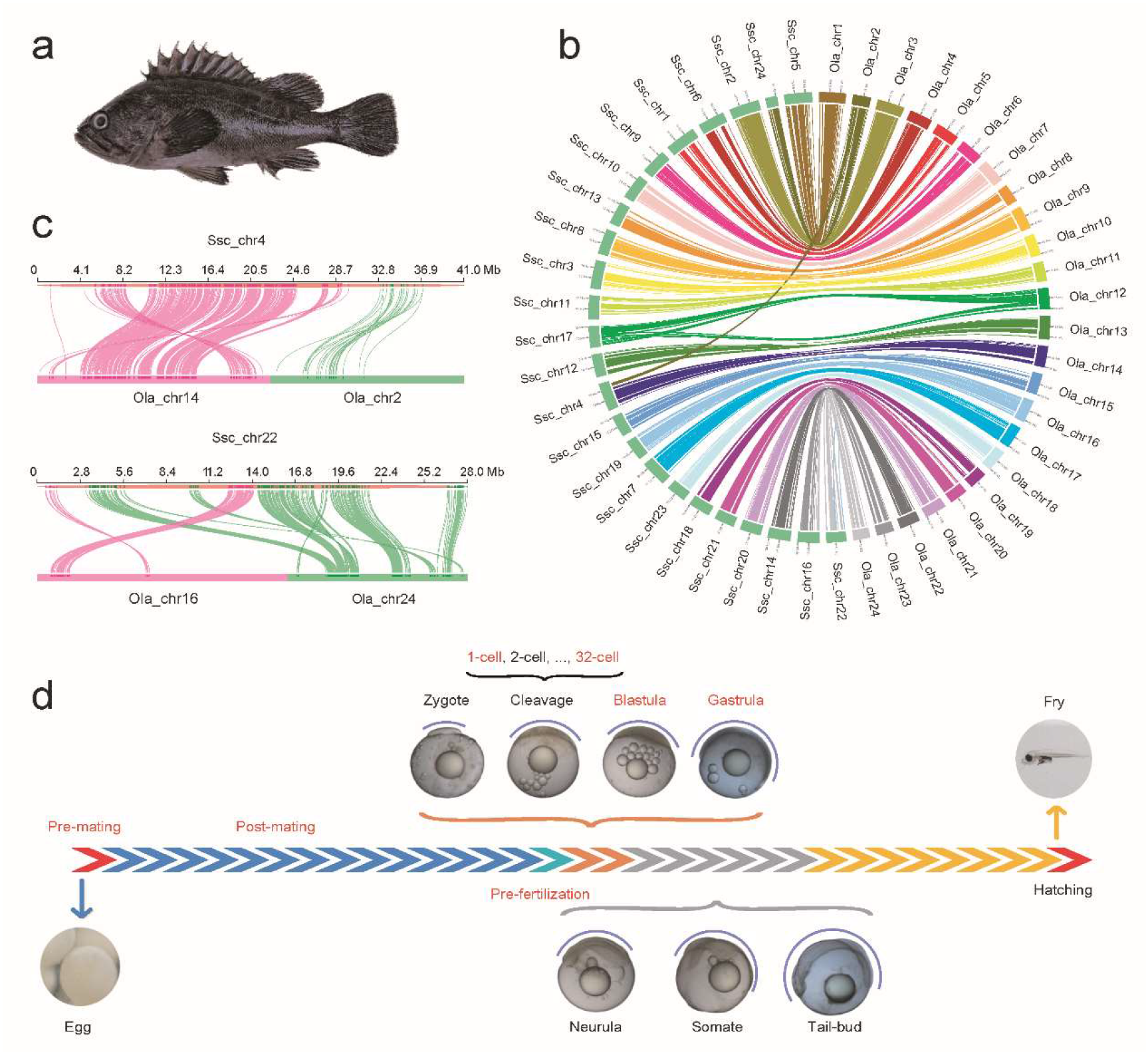
Synteny analysis and overview of rockfish reproduction. (a) Adult black rockfish (*Sebastes schlegelii*). (b) Synteny of rockfish and medaka genomes. (c) Recombination of chromosomes could only be observed in chr4 and chr22 of rockfish. (d) Schematic representation of rockfish reproduction and transcriptome sampling in the present study. Reproduction includes mating, pre-fertilization (six months; indicated in blue), embryo and early larvae development (hours; indicated in orange. Includes zygote, cleavage stage embryo, blastula, and gastrula), organogenesis (days; indicated in grey. Includes neurula, somite, and tail-bud), gestation (more than a month; indicated in yellow), and hatching. Purple marks the embryo proper. We sampled seven time points, highlighted in red font.

## Results

### Rockfish genome assembly and annotation

A critical first step in our effort to understand rockfish reproduction is the generation of an underlying high-quality genome assembly. We assembled the genome of a male rockfish (2n=48) by combining 57.3 Gb (~66×, genome estimation 868Mb based on *k*-mer analysis; Fig. S1, Tables S1) long PacBio reads and 114.6 Gb (~132×) short BGISEQ-500 reads (Methods, Tables S2). The genome assembly was 811 Mb, with a contig N50 size of 3.85 Mb (Table S3). We anchored ~99.86% of the assembled sequences onto 24 chromosomes using Hi-C (high-through chromosome conformation capture) data (Fig. S2, Table S4). Finally, we identified a 35.4% repeat content (Fig. S3, Table S5) and 24,094 protein-coding genes in the genome (Table S6). The structure of rockfish genes is similar to proximal species (Fig. S4, Table S7) and 99.71% genes could be annotated by a least one public database (Fig. S5, Table S8). To evaluate genome assembly quality, we first mapped short reads back to the final assembly, revealing a 98.13% mapping rate (Table S9). Using BUSCO (Benchmarking Universal Single-Copy Orthologues)^13^, we estimated the coverage of core vertebrate genes to be 93.9% and 94.4% in the assembly and gene set, respectively (Table S10). Furthermore, we found a good collinearity between rockfish and medaka genomes (Fig. 1b), with the exception of rockfish chromosomes 4 and 22. Each of them aligned to two medaka chromosomes (Fig. 1c), indicative of chromosome fusions. These assessments reflect the high quality of our rockfish genome assembly. Based on 1761 single-copy orthologs, we constructed a phylogenetic tree of rockfish and 15 other fish species. The tree suggests that rockfish (order Scorpaeniformes; a viviparous species with female parental care) and three-spined stickleback (order Gasterosteiformes; an egg-laying species with male parental care) diverged from a common ancestor approximately 84.9 Mya (Fig. S6), which corresponds to the Cretaceous period.

### Identification of gene expression correlating with maternal preparation

Black rockfish reproduction spans about eight months (Fig. 1d). Copulation occurs in November and December, while fertilization occurs approximately six months later, in April. The sperm storage stage is crucial, allowing maternal preparation for embryo development and hatching. In viviparous teleost fishes the ovary acts as both the source of eggs and the site in which eggs and embryos develop. After fertilization, the embryos develop in the ovary until hatching. Organogenesis completes within one day and is followed by about 50 days of gestation, when the offspring need to receive nutrition from the mother^14^.

The transcriptional program from mating to birth is highly stage- and cell/tissue-specific. We generated the transcriptomes of 21 adult tissues (Table S11) and carried out weighted gene co-expression network analysis (WGCNA)^15^ to identify genes expressed in concert in particular tissue(s) (termed modules) (Fig. 2a). Of 28 modules, two significantly correlated (*P* < 0.01) with a single sample type: TM08 with the oocyte and TM07 with the ovarian wall. Moreover, the expression of the genes within these modules were high in their correlating tissue. We further looked into functions of the genes in the two modules. Genes in TM07 are associated with processes likely important for maternal preparation for embryo implantation (Table S12). These include cell adhesion (collagen), blood vessel formation (*sox7, nln, vash1*, and *angpt2b*), response to blood vessel expansion and contraction (*ednrb*), guanytate cyclase activity (*gucy1a2, gucy2f*), NO-sGc-cGMP biosynthesis (*gucy1a2*), and extracellular calcium-sensing (*casr*). Other interesting genes in TM07 include a homolog to the oxygen-binding protein neuroglobin (*ngb*)^16,17^, and genes associated with trophoblast invasion into the maternal decidua (*htra3*)^18^ and smooth muscle development (*col12a1b* and *trpc4a*)^19^. These genes are related to early-stage embryo development in mammals; especially the NO/cGMP signalling pathway which plays an essential role in insemination, pregnancy, and birth^20–23^. These data suggest that maternal preparation of the ovarian wall is critical for rockfish viviparity.

**Fig. 2.**
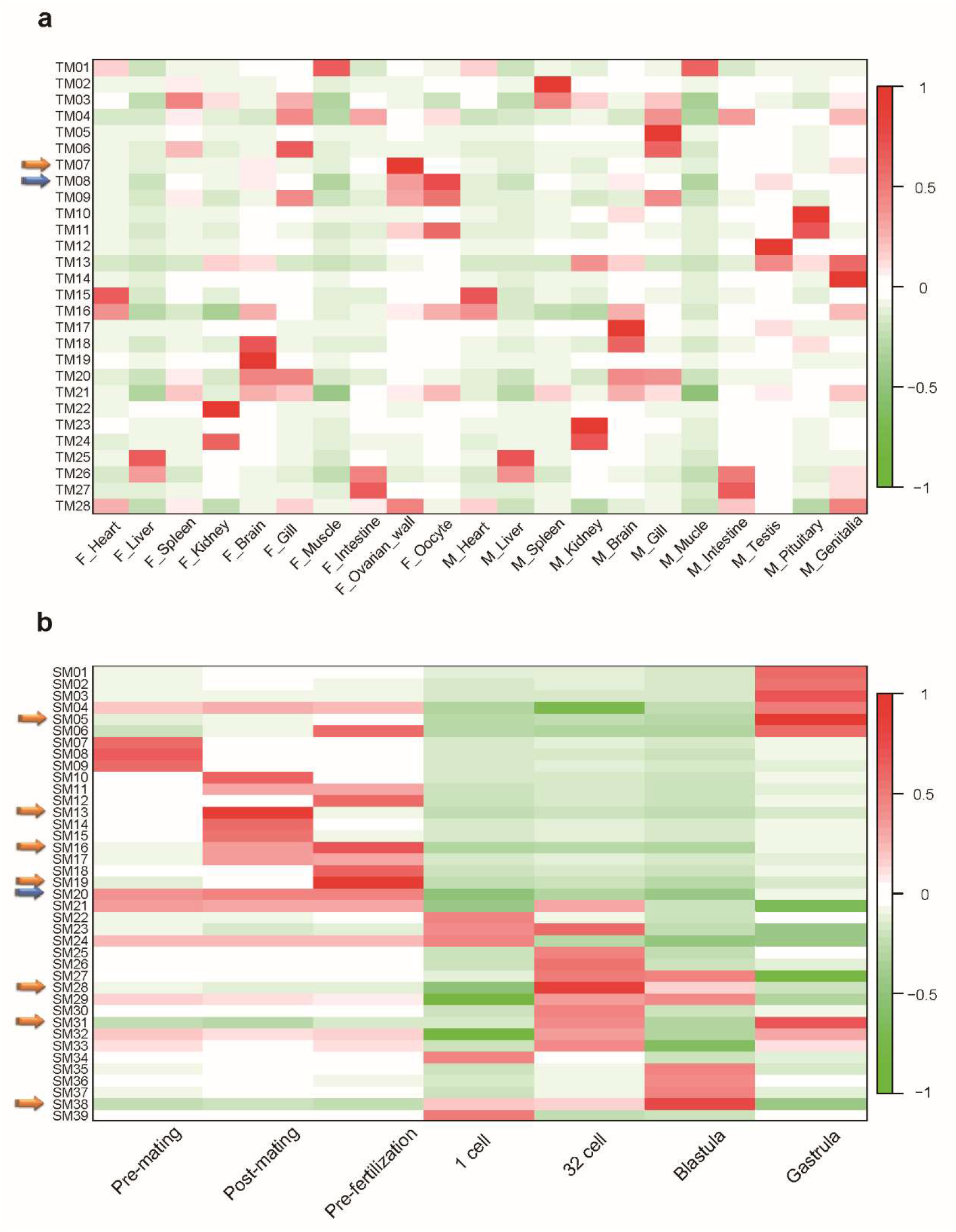
Identification of genes co-expressed in rockfish. (a) WGCNA co-expression modules were constructed by comparing (a) 21 tissues (b) seven reproduction time points. The *x*-axis shows sampled tissues in (a) and (b), with the prefix F_ for female and M_ for male samples, the *y*-axis WGCNA modules. For each module, the correlation value is indicated by a heat map ranging from −1 to 1. In total, 28 and 39 modules were identified in (a) and (b), respectively. Modules significantly correlated with a single tissue or stage are indicated by orange arrows; modules correlated with multiple tissues or stages by blue arrows.

We next obtained the transcriptomes of the pre-mating, post-mating, and pre-fertilization ovary; as well as the later 1-cell, 32-cell, blastula, and gastrula stage embryos (Table S13, Fig. 1d). In total, we sequenced 21 biological samples and carried out WGCNA (Fig. 2b). We identified more co-expressed genes in the pre-fertilization ovary (2,765 genes in SM16 and SM19) and gastrula embryos (4,998 genes in SM05 and SM31) compared to the post-mating ovary (141 genes in SM13) and the 32-cell (611 genes in SM28) and blastula embryos (343 genes in SM38). We did not detect any co-expressed modules in the pre-mating ovary and 1-cell embryo, indicating that these are relatively transcriptionally ‘dormant’ periods, whereas the pre-fertilization ovary and the gastrula embryos are more ‘active’. Furthermore, we identified a module (3,128 genes in SM20) with a large number of genes co-expressed by the ovary before fertilization (pre-mating, post-mating, and pre-fertilization). These genes were significantly enriched for several gene families (*P*-value < 0.05, Table 1, Table S14), including the zona pellucida (ZP) domain, prefoldin subunit, and DEAD/DEAH box helicase domain families. Zona pellucida, a component of the envelop surrounding fish eggs, is important for oogenesis, ovulation, fertilization, and embryogenesis. ZP genes constitute a species-restricted barrier for sperm at fertilization, act as a post-fertilization block to prevent polyspermy after gamete fusion, and contribute a hardened structure which protects the developing embryos until hatching^24^. Prefoldin subunit 1 (*pfd-1*) mutated animals with maternally contributed PFD-1 develop to the L4 larval stage and present with gonadogenesis defects which include aberrant distal tip cell migration^25^. Previous studies on DEAD-box proteins in model organisms have revealed their functions in the maintenance of gametogenesis^26^ and vascular endothelial growth^27^. DEAD-box proteins have an indispensable role in mammalian placental formation, which connect the developing embryo to the uterine wall and enables the delivery of oxygen and nutrients to the fetus and the return of metabolic wastes from the fetus to maternal circulation^28^.

**Table 1.** Protein family enrichment analysis of genes co-expressed by the ovary before fertilization. Enrichment analysis of genes in WGNCA module SM20 was carried out by searches against the STRING database online^29^.

| Pfam ID | Description | Gene number | *P*-value |
| --- | --- | --- | --- |
| PF00100 | Zona pellucida-like domain | 16 | 0.010 |
| PF02996 | Prefoldin subunit | 4 | 0.041 |
| PF00270 | DEAD/DEAH box helicase | 14 | 0.049 |

### Zona pellucida protein gene family in rockfish

To further dissect the role of ZP protein genes during viviparity, we annotated 22 ZP genes, aided by manual curation (Fig. S7, Table S15, S16), and performed *in situ* hybridization (ISH) and immunofluorescence (IF) to locate the position of sperm cells during post-mating and pre-fertilization. Sperm associated antigen 8 (*spag8*) ISH revealed strong mRNA staining of oocyte epithelium (Fig. 3a: 1-3). Furthermore, fluorescent staining of protamine 2 protein (PRM2) localized to the oocyte epithelium (Fig. 3a: 4-6). This indicates that sperm cells are in proximity of ZP proteins after mating. RNA-seq analysis showed that 19 out of 22 ZP genes are highly expressed before fertilization, while their expression after fertilization is relatively low (Fig. 3b, Table S17). We hypothesize that ZP proteins retain sperm after mating and that sperm is released for fertilization upon ZP protein degradation. To strengthen our hypothesis we performed chromatin accessibility profiling (ATAC-seq) during post-mating, pre-fertilization, and gastrula (Fig. S8). Consistently, a distinct difference in open chromatin regions adjacent to 12 ZP genes could be observed between maternal preparation (post-mating and pre-fertilization) and the gastrula stage (Fig. 3c, Fig. S9).

**Fig. 3.**
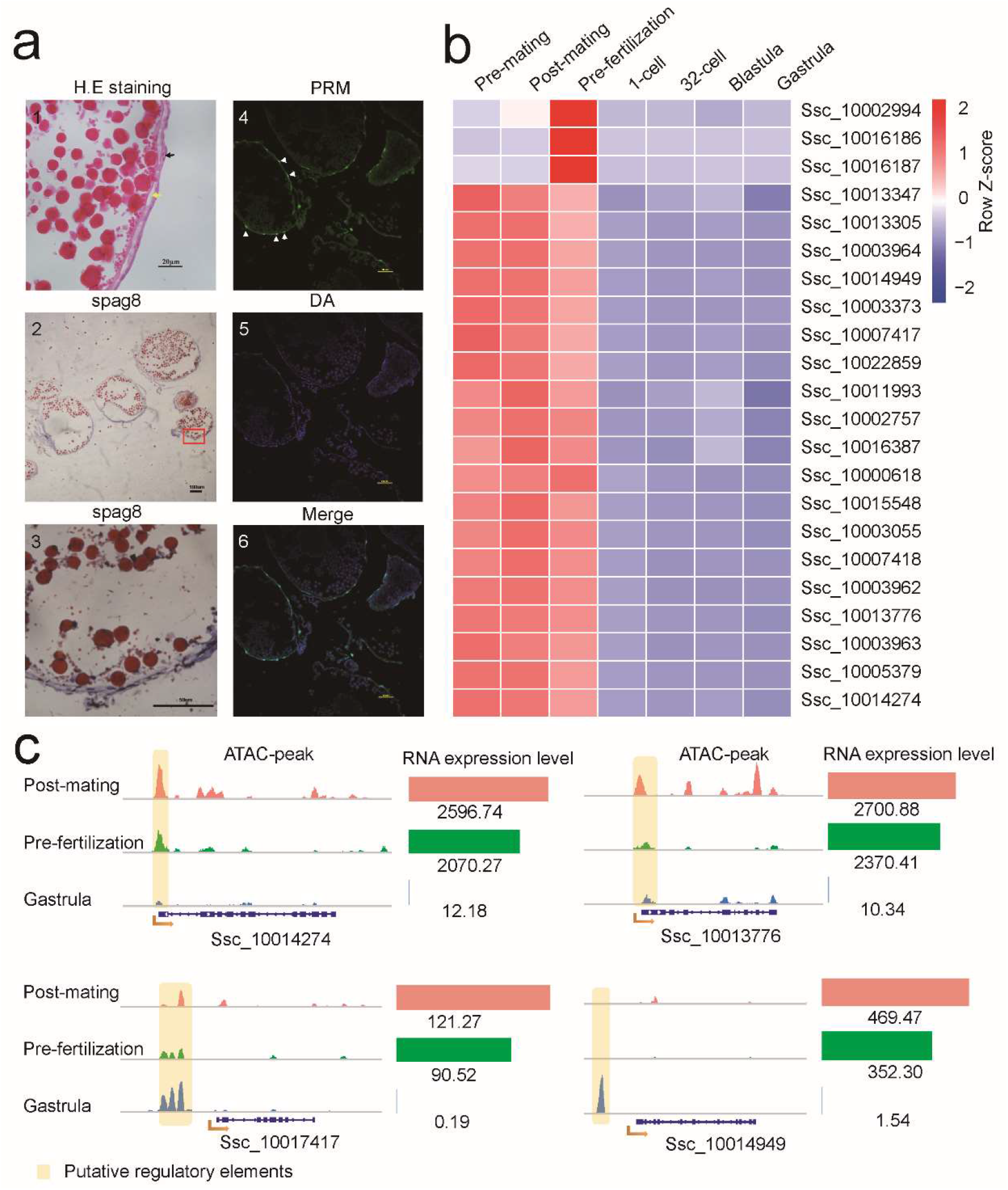
Zona pellucida protein gene family in rockfish. (a) Location of sperm and ZP on the surface of eggs. 1: Paraffin sections of an egg. H.E. staining. Scale bar=10μm. A number of spermatozoa surrounding the egg (black arrow) and the ZP region are indicated (yellow arrow); 2, 3: Expression of the spermatozoa marker *spag8* (DIG-labelled RNA) on the surface of eggs (blue region); 4, 5: Expression of the spermatozoa marker PRM2 (fluorescent antibody) on eggs. 6: Merged pictures. (b) Expression pattern of 22 ZP genes at seven time points. The log ratio expression is indicated in a heat map (−2 to 2; red: high, blue: low). (c) Signals of accessible chromatin and associated RNA expression levels of selected ZP genes from (b) in the post-mating and pre-fertilization ovary and gastrula stage of embryonic development. ATAC result of each gene is shown on the left, with peaks indicating accessible chromatin regions; gene expression levels (in TPM) on the right.

### Astacin metalloproteinase family in rockfish

After fertilization, polymerized and cross-linked ZP proteins are digested by hatching enzymes^30^, a subfamily of astacin metalloproteinases which lyse the chorion surrounding the egg, leading to hatching of embryos^31^. Astacin metalloproteinase can be classified into five subfamilies: hatching enzyme, HCE1-like, HCE2-like, patristacin/astacin, and nephrosin (Fig. 4a). We identified 26 astacin family genes, with two expanded subfamilies of hatching enzyme and HCE1-like in rockfish (Table S18, S19). These are distinct to the patristacin/astacin subfamily expansion in seahorse which play a role in brood pouch development and/or hatching of embryos within the brood pouch prior to parturition^32^. To examine the expression patterns of rockfish astacin genes we performed qRT–PCR of five time-points: pre-mating, post-mating, pre-fertilization, gastrula, and pre-hatching. Two of twelve HCE1-like genes (Ssc_10005765 and Ssc_10001812) were highly expressed pre-fertilization, while two of the eight hatching enzyme genes (Ssc_10008384 and Ssc_10021724) were highly expressed pre-hatching (Fig. 4b). Thus, we propose that astacin family members play distinct roles in rockfish viviparity. HCE1-like proteins play a role in releasing spermatozoa from the zona pellucida at the pre-fertilization stage, while hatching enzymes are responsible for freeing the embryo from the chorion at the pre-hatching stage.

**Fig. 4.**
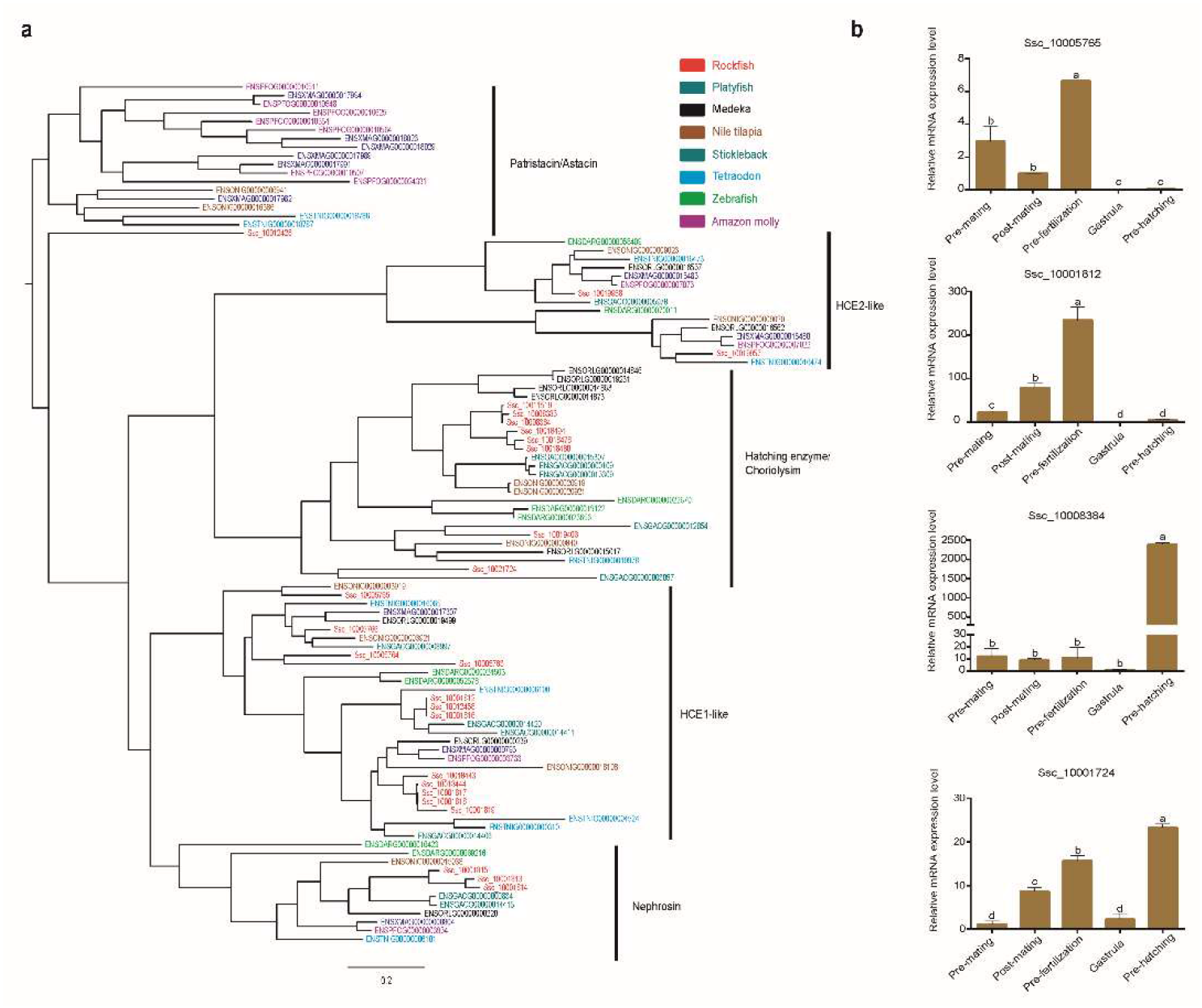
Assessment of the astacin metalloproteinase family. (a) Phylogenetic tree of astacin family genes in eight fish species. In rockfish eight genes were found in the hatching enzyme subfamily and 12 genes in the HCE1-like subfamily (b) the expression profiles of four hatching enzyme genes in rockfish, quantified by qRT–PCR. Columns with different letters (a–b) represents significantly different expression (*P* < 0.05, one-way ANOVA). Data are mean±s.e.m. of three biological replicates.

## Discussion

In this study, we present a high-quality genome assembly of the rockfish *Sebastes schlegelii*, a viviparous fish in the teleost order Scorpaeniformes. We generated transcriptomes of different tissues and different developmental time points, to reveal gene expression patterns related to viviparity. The reproduction of viviparous rockfish is well-established, however the genetic changes associated with the long period between copulation and fertilization (six months, including maternal preparation and sperm storage), embryo development, and hatching has hitherto remained unknown. We found that ZP proteins and HCE-1 like proteins likely play important roles during sperm storage and release, respectively (Fig. 5). We propose that ZP proteins retain and stabilize sperm in lamellae epithelium prior to fertilization – a six-month period in rockfish. In contrast, HCE1-like proteins digest the zona pellucida and contribute to sperm release. In the maternal preparation stage, the preparation for ‘embryo implantation’, we found expression of genes related to cell adhesion, trophoblast invasion, calcium-sensing receptors, the NO-sGc-cGMP signaling pathway, and blood vessel function. Significant differences in the internal circulation system of fish embryos has previously been reported^33^, believed to due to the high demand for oxygen associated with parental care. In rockfish we not only observed high expression of vasodilatation and angiogenesis-related genes, but also genes associated with the growth of blood vessels in the ovarian wall during reproduction. For example, a homolog of the oxygen-binding neuroglobin was highly expressed by the ovarian wall. In mammals, the function of the uterus is to transfer oxygen and nutrients to the growing embryo. The chemical messenger nitric oxide (NO) is a key regulator of fetoplacental circulation in mammals^23^ and also induces oocyte maturation in the zebrafish via the NO-sGc-cGMP pathway^20^. In the rockfish ovarian wall we observed high expression of nitric oxide synthase 1 adaptor protein (*nos1ap;* also known as *CAPON*), a regulator of NO bioavailability^34^. Rockfish embryos not only receive parental care, but also parental nutrition^14^. We provide evidence, at the transcriptome level, for maternal preparation for embryo implantation and the formation of an interface between mother and offspring (embryo). We hypothesize that the ovarian wall of the rockfish functions similar to the mammalian uterus.

**Fig. 5.**
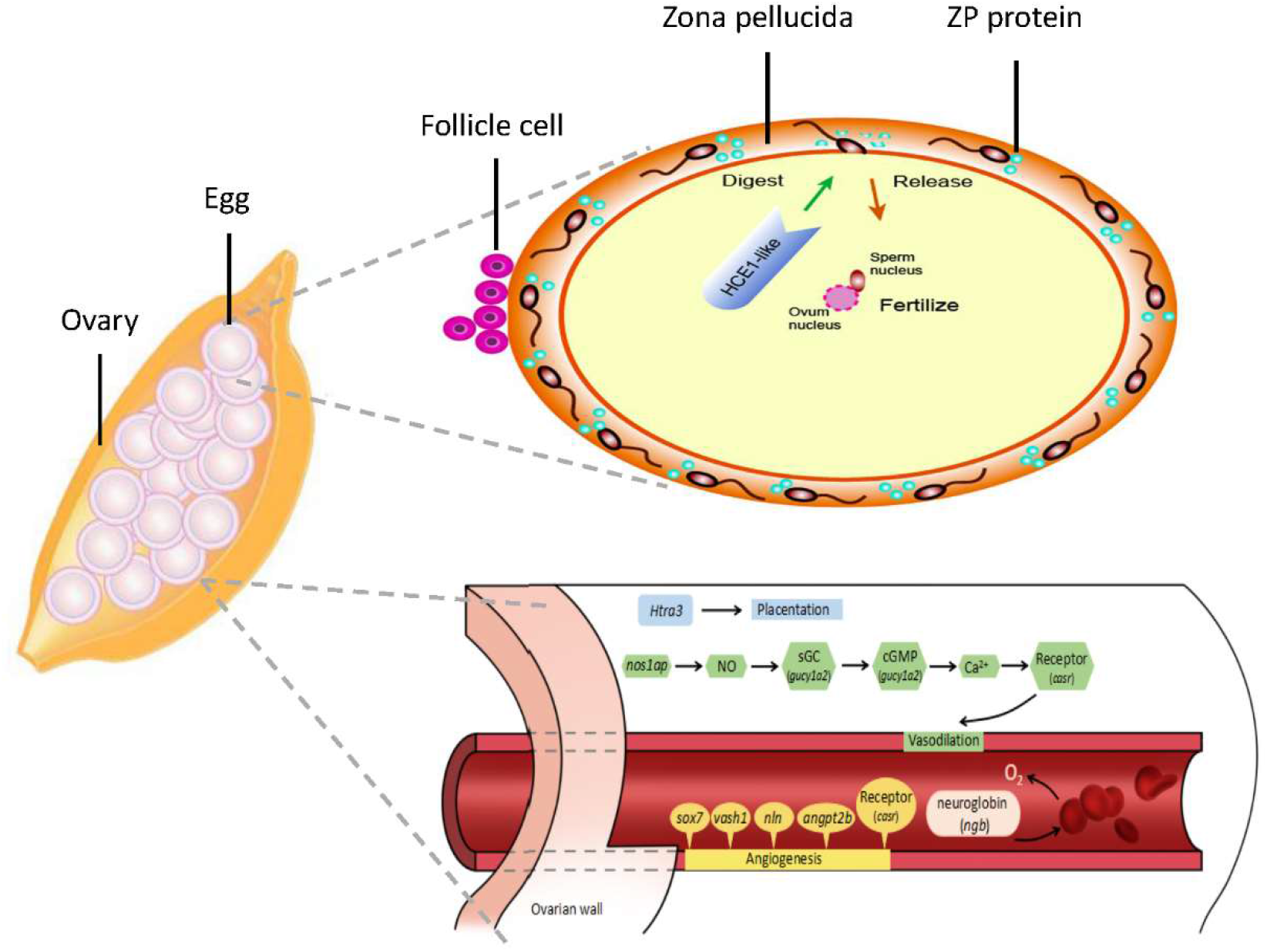
Proposed model of viviparous reproduction in rockfish. Sperm is retained in the zona pellucida (ZP) region of eggs and released for fertilization upon degradation of ZP proteins. The rockfish ovarian wall is analogous to the mammalian uterus.

In conclusion, we have generated and analysed a high-quality genomic data set for black rockfish to generate a model of its reproduction – the first such model of fish viviparity. This model, along with the candidate genes identified in our study, will greatly facilitate future studies on the evolution of fish, as well as vertebrate life history strategies.

## Methods

### Ethics statement

This study was approved by the Animal Care and Use Committee of the Centre for Applied Aquatic Genomics at the Ocean University of China.

### Samples

Black rockfish (*Sebastes schlegelii*) were obtained from the Zhucha Island (Qingdao, Shandong, China). A three-year-old male adult (weight 665g) was used for genome sequencing. Samples used for RNA- and ATAC-sequencing were collected from November 2017 to March 2018. Twelve healthy three-year-old fish (six males and six females) were randomly selected for sampling of heart, liver, spleen, kidney, pituitary, brain, intestine, gill, muscle, genitalia (testis or ovary), ovarian wall, and oocytes in November 2017. The ovary of the female at different developmental stages (pre-mating, post-mating, and pre-fertilization) were collected on November 2017, February 2018, and March 2018, respectively. Embryos (20-30 each) at various developmental stages including 1-cell, 32-cell, blastula, gastrula and pre-hatching were collected.

### Sequencing, assembly and evaluation of rockfish genome

The black rockfish genome was assembled using CANU v1.2 software^35^ with the parameters ‘MhapSensitivity=high corMinCoverage=0 minReadLength=500 genomeSize=868m errorRate=0.04’ in three steps: correct, trim, and assemble. Though the assembly had been corrected by CANU, a strict error-correcting procedure was performed: firstly, the draft genome was corrected using PacBio long reads, secondly, the assembly was further corrected using Pilon v1.22^36^ and 98.1G (114.6X) WGS short reads.

To generate a chromosomal-level assembly of the genome, we took advantage of sequencing data from a Hi-C library^37^. We performed quality control of Hi-C raw data using HiC-Pro (v2.8.0)^38^. We used bowtie2 (v2.2.5)^39^ to compare raw reads to the draft assembled genome, filtering out low-quality reads to build raw inter/intra-chromosomal contact maps. Our final data set was 41.75 Gb (48.1×), accounting for 54.59% of the total Hi-C sequencing data. We next used Juicer (v1.5)^40^, an open-source tool to analyse Hi-C datasets, and a 3D *de novo* assembly (3d-dna, v170123) pipeline to scaffold spotted rockfish genome to chromosomes. We further conducted whole-genome alignment between the rockfish genome and the published medaka genome using LASTZ (v1.10)^41^ to compare consistency between these two genomes.

### Completeness assessment of genome assemblies

To evaluate the consistence and intergrity of genome assembly, short-insert library reads were used to map with the assembled genome using BWA^42^ to generate mapping ratio statistics. BUSCO (Benchmarking Universal Single-Copy Orthologues)^13^ provides quantitative measures for the assessment of genome assembly completeness, based on evolutionarily-informed expectations of gene content from near-universal single-copy orthologs.

### Repeat and gene annotation

We constructed a transposable element (TE) library of rockfish genome using a combination of homology-based and *de novo* approaches. For the *ab-initio* method, we used RepeatModeler and LTR_FINDER^43^ to build the rockfish custom repeat database. In homology-based method, we used the Repbase database^44^ to identify repeats with RepeatMasker and RepeatProteinMask (http://www.repeatmasker.org).

The annotation strategy of *Rockfish* protein-coding genes was to integrate *de novo* prediction and evidence based including homology and transcriptome data. Protein sequences of zebrafish (*Danio rerio*, fugu (*Takifugu rubripes*), spotted green pufferfish (*Tetraodon nigroviridis*), stickleback (*Gasterosteus aculeatus*), large yellow croaker (*Larimichthys crocea*), tongue sole (*Cynoglossus semilaevis*), tilapia (*Oreochromis niloticus*), medaka (*Oryzias latipes*), and Amazon molly (*Poecilia formosa*) were downloaded from Ensembl (http://ensemblgenomes.org/) or NCBI (https://www.ncbi.nlm.nih.gov/genome/). These sequences were aligned to the rockfish genome with tBLASTn (*E*-value ⩽ 10^−5)^ and GeneWise^45^ was used to generated gene structure based on the alignment. The transcriptome data (including 11 tissues in Method Fish section) were assembled by Trinity^46^ and mapped to rockfish genome by BLAT ^47^. The *de novo* prediction of rockfish was carried out with Augustus^48^. All evidences of gene model were integrated with GLEAN^49^.

### RNA-seq and transcriptome data processing

In total, 71 RNA-seq libraries, 58 of which from 21 tissues, and 29 from seven different developmental stages (pre-mating, post-mating, pre-fertilization, 1-cell, 32-cell, blastula, gastrula) with three biological replicates. These libraries were sequenced 100 bp at each end using BGI-seq 500 platform. The transcriptomic data were mapped to the rockfish genome and the expression level of genes was calculated using Salmon^50^ with default parameters. The genes of interest were subsequently visualized by Heatmap package. In order to the investigate the regulatory network, we used the gene co-expression network constructed by the method of Weighted Gene Co-Expression Network Analysis (WGCNA)^15^.

### ATAC experiment and data processing

We treated the samples at pre-, post-mating and gastrula stage and build the ATAC-seq libraries with a protocol modified from previous report^51^. All ATAC-seq libraries were sequenced using BGI-SEQ500. After filtering low quality data, duplication and removing chondriosome DNA, the clean reads were aligned to the genomes using Bowtie2 (version 2.2.2)^52^. All the ATAC-seq peaks were called by MACS^53^ with the parameters ‘nolambda –nomodel’.

### Prediction and clustering of ZP and Astacin metalloproteinase

The previously reported ZP and astacin metalloproteinase in zebrafish and medaka^32,54^ were manually verified, and subsequently used as baits to predict the target proteins in rockfish. The candidates were filtered with domain information obtained with SMART^55^ and confirmed by blast against NR database. Furthermore, multiple sequence alignment of ZP and astacin metalloproteinase was respectively carried out using MUSCLE^56^, with both predicted and reported proteins in rockfish, medaka, zebrafish, tilapia, Amazon molly, platyfish, tetraodon and stickleback^32,54^. Phylogenetic trees were subsequently constructed based on the alignment results with Fasttree^57^ by Maximum Likelihood.

### RNA isolation, cDNA synthesis and quantitative real-time RT-PCR

Total RNA was extracted using TRIzol (Invitrogen, Carlsbad CA, USA) and complementary DNA (cDNA) was synthesized using the Reverse Transcriptase M-MLV Kit (TaKaRa). All quantitative real-time RT-PCR experiments were performed in triplicate on a Light-Cycler Roche 480 instrument (Roche Applied Science, Mannheim, Germany), using primers against hatching enzyme genes and the housekeeping gene ribosomal protein L17 (*rpl17*) (primers shown in **Table S20**). The relative expression of each hatching enzyme gene was calculated using the comparative 2^−ΔΔCt^ method^58^. The results were expressed as mean ± standard error of the mean (s.e.m.). Data was evaluated using a one-way ANOVA.

### *In situ* hybridization

In *situ* hybridization of ovaries were conducted as described^32^. Antisense mRNA probes of *spag8*, a marker of spermatozoa, were synthesized using a DIG RNA Labelling Mix (Indianapolis, IN, USA). A pair of primers (*Ssc-spag8*-Fw/Rv) was used for probe synthesis (**Table S20**). The results were photographed by AZ100 (Nikon, Tokyo, Japan).

### Immunofluorescence

Ovaries of fishes were removed, cleaned of excess fat, fixed in 4% formaldehyde solution overnight at 4 °C, and dehydrated to 100% for histological analysis. Ovaries were serially-sectioned at 7 μm on a RM2016 Paraffin slicer onto glass slides and washed in 1x Phosphate-Buffered Saline (PBS) containing 0.1% Triton-X (CalBiochem). Tissue sections were incubated in blocking buffer (3% Goat Serum; Sigma), 1% Bovine Serum Albumin (Sigma), and 0.5% Tween-20 (Fisher Scientific) in 1X PBS], and incubated with rabbit primary antibodies against the spermatozoa marker protamine 2 protein (PRM2; Uscn Life Science Inc., PAH307Hu01). A secondary FITC AffiniPure Goat Anti-Rabbit IgG antibody was used to enable green fluorescence detection. Sections were stained with DAPI to visualize nuclei and analysed on a laser scanning confocal microscope.

## Acknowledgements

We give our special thanks to He Zhang from Sichuan Fine Art Institute, who kindly provided help with drawing fig. 5. This study was financially supported by grants from the fundamental Research Funds for Central Universities (No. 201822026) and National Key R&D Program of China (2018YFD0900101, 2018YFD0901205 and 2018YFD0900301).

## Author contributions

Yan He, Guangyi Fan, Quanqi Zhang, Huanming Yang, Jian Wang and Jie Qi conceived the study; Yan He, Yue Chang, Lisui Bao, Mengjun Yu, He Zhang and Weihao Song interpreted the data. Yan He, Rui Li, Yating Qin, Yilin Wang, Longqi Liu, Jingjing Niu, Xuemei Li, Xiangfu Kong, Meiting Peng, Minmin Sun, Mengya Wang, Jiangbo Qu, Xiaobing Liu, Jingxiang Liu, Xiaolong Wu, Xi Zhao and Xuliang Wang performed the experiments. Haiyang Yu, Xubo Wang, Jie Cheng, Xuangang Wang, Zhigang Wang, Yaolei Zhang, Jiao Guo, Yang Liu and Kaiqiang Liu prepared the material; Yan He, Yue Chang, Lisui Bao, Mengjun Yu, Guangyi Fan and Jie Qi drafted the manuscript. Xin Liu, Guangyi Fan, Inge Seim, Yue Chang, Mengjun Yu, Yan He, Quanqi Zhang, Simon Ming-Yuen Lee, Xun Xu and Jie Qi contributed to the final manuscript editing.

## Data Accession

The project has been deposited at CNSA(CNGB Nucleotide Sequence Archive) under the accession ID CNP0000222. The assembled genome can be obtained by assembly ID CNA0000824.

## References

1. Blackburn, D. G. in Encyclopedia of reproduction Vol. 3 (ed E. Knobil and J. D. Neill) 994–1003 (Academic Press, 1999).

2. Helmstetter, A. J. et al. Viviparity stimulates diversification in an order of fish. Nature communications 7, 11271 (2016).

3. Wootton, R. J. & Smith, C. in Reproductive biology of teleost fishes Ch. 9, 252–279 (John Wiley & Sons, Ltd, 2014).

4. Wourms, J. P. Viviparity: the maternal-fetal relationship in fishes. American Zoologist 21, 473–515 (1981).

5. Nelson, J. S. in Fishes of the World. 4th (ed Joseph S. Nelson) 1–625 (John Wiley & Sons, Inc., 2006).

6. Haynes, J. L. Standardized classification of poeciliid development for life-history studies. Copeia, 147-154 (1995).

7. Lucinda, P. H. Family Poeciliidae. Check list of the freshwater fishes of South and Central America, 555–581 (2003).

8. Thibault, R. E. & Schultz, R. J. Reproductive adaptations among viviparous fishes (Cyprinodontiformes: Poeciliidae). Evolution 32, 320–333 (1978).

9. Schartl, M. et al. The genome of the platyfish, Xiphophorus maculatus, provides insights into evolutionary adaptation and several complex traits. Nature genetics 45, 567–572 (2013).

10. Omoto, N. et al. Gonadal sex differentiation and effect of rearing temperature on sex ratio in black rockfish (Sebastes schlegeli). Ichthyological research 57, 133–138 (2010).

11. Kusakari, M. Mariculture of kurosoi, Sebastes schlegeli. Environmental Biology of Fishes 30, 245–251 (1991).

12. Mori, H., Nakagawa, M., Soyano, K. & Koya, Y. Annual reproductive cycle of black rockfish Sebastes schlegeli in captivity. Fisheries science 69, 910–923 (2003).

13. Simão, F. A., Waterhouse, R. M., Ioannidis, P., Kriventseva, E. V. & Zdobnov, E. M. BUSCO: assessing genome assembly and annotation completeness with single-copy orthologs. Bioinformatics 31, 3210–3212 (2015).

14. Boehlert, G. W., Kusakari, M. & Shimizu, M. Energetics during embryonic development in kurosoi, Sebastes schlegeli Hilgendorf. Journal of experimental marine biology and ecology 101, 239–256 (1986).

15. Langfelder, P. & Horvath, S. WGCNA: an R package for weighted correlation network analysis. BMC bioinformatics 9, 559 (2008).

16. Pesce, A. et al. Neuroglobin and cytoglobin: Fresh blood for the vertebrate globin family. EMB Oreports 3, 1146–1151 (2002).

17. Brunori, M. et al. Neuroglobin, nitric oxide, and oxygen: functional pathways and conformational changes. Proceedings of the National Academy of Sciences 102, 848–38488 (2005).

18. Singh, H., Endo, Y. & Nie, G. Decidual HtrA3 negatively regulates trophoblast invasion during human placentation. Human reproduction 26, 748–757 (2011).

19. Tsvilovskyy, V. V. et al. Deletion of TRPC4 and TRPC6 in mice impairs smooth muscle contraction and intestinal motility in vivo. Gastroenterology 137, 1415–1424 (2009).

20. Li, J., Zhou, W., Wang, Y. & Niu, C. The dual role of cGMP in oocyte maturation of zebrafish. Biochemical and biophysical research communications 499, 998–1003 (2018).

21. Sladek, S. M., Magness, R. R. & Conrad, K. P. Nitric oxide and pregnancy. American Journal of Physiology-Regulatory, Integrative and Comparative Physiology 272, R441–R463 (1997).

22. Dantas, B. P. et al. Vasorelaxation induced by a new naphthoquinone-oxime is mediated by NO-sGC-cGMP pathway. Molecules 19, 9773–9785 (2014).

23. Tropea, T. et al. Nitrite mediated vasorelaxation in human chorionic plate vessels is enhanced by hypoxia and dependent on the NO-sGC-cGMP pathway. Nitric Oxide 80, 82–88 (2018).

24. Ahn, D.-H. et al. Draft genome of the Antarctic dragonfish, Parachaenichthys charcoti. GigaScience 6, 1–6 (2017).

25. Lundin, V. F., Srayko, M., Hyman, A. A. & Leroux, M. R. Efficient chaperone-mediated tubulin biogenesis is essential for cell division and cell migration in C. elegans. Developmental biology 313, 320–334 (2008).

26. Kotov, A., Akulenko, N., Kibanov, M. & Olenina, L. DEAD-Box RNA helicases in animal gametogenesis. Molecular Biology 48, 16–28 (2014).

27. de Vries, S. et al. Identification of DDX6 as a cellular modulator of VEGF expression under hypoxia. Journal of biological chemistry 288, 5815–5827 (2013).

28. Chen, C.-Y. et al. Targeted inactivation of murine Ddx3x: essential roles of Ddx3x in placentation and embryogenesis. Human molecular genetics 25, 2905–2922 (2016).

29. Szklarczyk, D. et al. The STRING database in 2017: quality-controlled protein–protein association networks, made broadly accessible. Nucleic Acids Research 45, D362–D368 (2017).

30. Litscher, E. S. & Wassarman, P. M. in Extracellular Matrix and Egg Coats (eds E. S. Litscher & P. M. Wassarman) Ch. 9, 2–488 (Elsevier, 2018).

31. Kawaguchi, M. et al. Evolution of teleostean hatching enzyme genes and their paralogous genes. Development Genes & Evolution 216, 769–784 (2006).

32. Lin, Q. et al. The seahorse genome and the evolution of its specialized morphology. Nature 540, 395–399 (2016).

33. Ho, D. H. Transgenerational epigenetics: The role of maternal effects in cardiovascular development. Integrative and Comparative Biology 54, 43–51 (2014).

34. Candemir, E. et al. Interaction of NOS1AP with the NOS-I PDZ domain: implications for schizophrenia-related alterations in dendritic morphology. European Neuropsychopharmacology 26, 741–755 (2016).

35. Koren, S. et al. Canu: scalable and accurate long-read assembly via adaptive k-mer weighting and repeat separation. Genome research 27, 722–736 (2017).

36. Walker, B. J. et al. Pilon: An Integrated Tool for Comprehensive Microbial Variant Detection and Genome Assembly Improvement. Plos One 9, e112963 (2014).

37. Burton, J. N. et al. Chromosome-scale scaffolding of de novo genome assemblies based on chromatin interactions. Nature biotechnology 31, 1119–1125 (2013).

38. Servant, N. et al. HiC-Pro: an optimized and flexible pipeline for Hi-C data processing. Genome biology 16, 259 (2015).

39. Langmead, B., Trapnell, C., Pop, M. & Salzberg, S. L. Ultrafast and memory-efficient alignment of short DNA sequences to the human genome. Genome biology 10, R25 (2009).

40. Durand, N. C. et al. Juicer provides a one-click system for analyzing loop-resolution Hi-C experiments. Cell systems 3, 95–98 (2016).

41. Harris, R. S. Improved Pairwise Alignmnet of Genomic DNA. PhD thesis, Pennsylvania State Univ. (2007).

42. Li, H. & Durbin, R. Fast and accurate short read alignment with Burrows–Wheeler transform. bioinformatics 25, 1754–1760 (2009).

43. Xu, Z. & Wang, H. LTR_FINDER: an efficient tool for the prediction of full-length LTR retrotransposons. Nucleic acids research 35, W265–W268 (2007).

44. Bao, W., Kojima, K. K. & Kohany, O. Repbase Update, a database of repetitive elements in eukaryotic genomes. Mobile DNA 6, 11 (2015).

45. Birney, E., Clamp, M. & Durbin, R. GeneWise and genomewise. Genome research 14, 988–995 (2004).

46. Grabherr, M. G. et al. Trinity: reconstructing a full-length transcriptome without a genome from RNA-Seq data. Nature biotechnology 29, 644–652 (2011).

47. Kent, W. J. BLAT—the BLAST-like alignment tool. Genome research 12, 656–664 (2002).

48. Stanke, M. & Waack, S. Gene prediction with a hidden Markov model and a new intron submodel. Bioinformatics 19, ii215–ii225 (2003).

49. Elsik, C. G. et al. Creating a honey bee consensus gene set. Genome biology 8, R13 (2007).

50. Patro, R., Duggal, G., Love, M. I., Irizarry, R. A. & Kingsford, C. Salmon provides fast and bias-aware quantification of transcript expression. Nature methods 14, 417–419 (2017).

51. Wu, J. et al. The landscape of accessible chromatin in mammalian preimplantation embryos. Nature 534, 652–657 (2016).

52. Langmead, B. & Salzberg, S. L. Fast gapped-read alignment with Bowtie 2. Nature methods 9, 357–359 (2012).

53. Zhang, Y. et al. Model-based analysis of ChIP-Seq (MACS). Genome biology 9, R137 (2008).

54. Wu, T. et al. Bioinformatic analyses of zona pellucida genes in vertebrates and their expression in Nile tilapia. Fish physiology and biochemistry 44, 435–449 (2018).

55. Schultz, J., Copley, R. R., Doerks, T., Ponting, C. P. & Bork, P. SMART: a web-based tool for the study of genetically mobile domains. Nucleic acids research 28, 231–234 (2000).

56. Edgar, R. C. MUSCLE: multiple sequence alignment with high accuracy and high throughput. Nucleic acids research 32, 1792–1797 (2004).

57. Price, M. N., Dehal, P. S. & Arkin, A. P. FastTree: computing large minimum evolution trees with profiles instead of a distance matrix. Molecular biology and evolution 26, 1641–1650 (2009).

58. Livak, K. J. & Schmittgen, T. D. Analysis of relative gene expression data using real-time quantitative PCR and the 2− ΔΔCT method. methods 25, 402–408 (2001).

