## supplementary files for "A chromosome-level genome of black rockfish, *Sebastes schlegelii*, provides insights into the evolution of live birth"

### Chromosome level genome sequence of *Sebastes schlegelii* reveals the regulatory elements and genes related to reproduction of viviparous fish

This PDF file includes:

- Supplementary Text
- Figs. S1to S15
- Table S1 to S10,S14,S15,S17-S20
- References(1-31)

Other Supplementary Materials for this manuscript include the following:

- Table S11-S13, Table S16

### Supplementary Text

#### Sample preparation and genome sequencing

All experimental black rockfish (*Sebastes schlegelii*) and embryos were obtained from the Huangdao Aquaculture Market (Qingdao, Shandong Province, China). A three-year-old male adult (weight 665g) was used for genome sequencing and DNA extracted from blood tissue. Fishes were collected every other month from November 2017 to March 2018. Twelve healthy 3-year-old fish (six males and six females) were randomly selected for heart, liver, spleen, kidney, pituitary, brain, intestine, gill, muscle, ovarian wall, genitalia and gonad (i.e., ovary and testis) sampling in Novermber 2017. The gonads of the fish at different developmental stages (pre-mating, post-mating, pre-fertilization) were collected. One side of the gonad was collected for RNA extraction, while the other was fixed in 4% PFA solution for histological analysis, *in situ hybridization* (ISH) and immunofluorescence. Every 20–30 embryos at various developmental stages (1 cell, 32 cell, blastula, gastrula, neurula, somatic stage, tail-bud stage, 24hpf (hours post fertilization) and 48hpf were collected.

Total RNA from the above collected samples was extracted by TRIzol Regent (Invitrogen, Carlsbad CA, USA) according to the manufacturer's instructions, and treated with RNase-free DNase 1 (TaKaRa, Dalian, China) to degrade residual DNA. The cDNA was transcribed with 1μg of total RNA using the Reverse Transcriptase M-MLV Kit (TaKaRa) following the manufacture's protocols.

#### Estimate the Genome Size using K-mer spectrum

The genome size of rockfish was estimated based on K-mer spectrum. Given the K-mer frequency obeys a Poisson distribution, when the coverage is sufficient, genome size can be estimated by the formula: Genome Size = K*num*/K*depth*. Where K-num is the number of K-mers, and K*depth* is the expected depth of K-mers. In this study, 98.1G (114.6-fold) PE150 short reads sequencing with Illumina platform were used for K-mer analysis and the K-num is 49,484,519,640 based on 17-mers and K-mer depth is 57. Therefore, we estimated the genome size is around 868 Mb (**Fig. S1, Table S1**).

#### Genome assembly

The rockfish genome was assembled using CANU[1]software with parameters "MhapSensitivity=high corMinCoverage=0 minReadLength=500 genomeSize=868m errorRate=0.04" using 57.3G(66-fold) PacBio data (**Table S2**)in three steps: Correct, Trim and Assemble. Even if the assembly had been corrected by CANU, A strict error-correcting procedure was launched in two steps: firstly, the draft genome was corrected by Arrow using PacBio long reads, Then, the assembly continue to be corrected by Pilon v1.22[2] using 98.1G(114.6-fold) PE150 short reads (**Table S2**). The final rockfish draft genome is 811M and Contig N50 is 3.85Mbp (**Table S3**).

To further generate a chromosomal-level assembly of the genome, we took advantage of sequencing data from the Hi-C[3] library. We performed quality control of Hi-C raw data using HiC-Pro (v. 2.8.0)[4]. First, we used bowtie2 (v. 2.2.5)[5] to compare the raw reads to the draft assembled sequence, and then low-quality reads were filtered out to build raw inter / intra-chromosomal contact maps. Our final valid data set was 41.75Gb (48.1×), accounting for 54.59% of the total Hi-C sequencing data. We then used Juicer (v. 1.5)[6], an open-source tool for analyzing Hi-C datasets, and 3D *de novo* assembly (3d-dna, v. 170123) pipeline, to scaffold the rockfish genome to 24 pseudochromosomes with length ranging from 16.58Mb to 43.93Mb (**Table S4, Fig. S2**). The total length of pseudochromosomes consisted of 99.73% of all genome sequences. We further conducted whole genome alignment between the rockfish genome and the published medaka genome using LASTZ (v. 1.10) [7] to compare consistency between these two genomes. The 24 pseudochromosomes we identified in black rockfish genome aligned exactly against the 24 chromosomes of the medaka genome (**Fig. 1a**).

**Transposable element analysis**

We constructed a transposable element (TE) library of rockfish genome using a combination of homology-based and *de novo* approaches. For the *ab-initio* method, we used RepeatModeler and LTR_FINDER[8] to build the rockfish specific repeat database. In homology-based method, we used known repeat library (Repbase) [9] to identify repeats with RepeatMasker (the parameter setting as '-a -nolow -no_is -norna -parallel 3 -e wublast --pvalue 0.0001') (http://www.repeatmasker.org) and RepeatProteinMask (the parameter setting as '-noLowSimple -pvalue 0.0001 -engine wublast') (http://www.repeatmasker.org). Tandem repeats finder (TRF)[10] was used to find tandem repeats. The rockfish harbors 35.39% of repeats sequence (**Table S5, Fig. S3**).

**Gene prediction and annotation**

The annotation strategy of rockfish protein-coding genes was to integrate *de novo* prediction and evidence based including homology and transcriptome data. Homology sequences of species including *Danio rerio*, *Takifugu rubripes*, *Tetraodon nigroviridis*, *Gasterosteus aculeatus*, *Larimichthys crocea*, *Cynoglossus semilaevis*, *Oreochromis niloticus*, *Oryzias latipes*, *Poecilia formosa* were downloaded from Ensemble (http://ensemblgenomes.org/ ) or NCBI (https://www.ncbi.nlm.nih.gov/genome/). The protein sequences of homology species were aligned to rockfish genome with TBLASTn (e-value $\leq 10^{-5)}$ and generated gene structure with GeneWise[11] (the parameter setting as '-genesf'). The transcriptome data (including liver, kidney, brain, muscle, ovarian wall, oocyte in female and liver, kidney, brain, muscle, testis in male) were assembled by Trinity [12] and mapped to rockfish genome by BLAT[13]. The *de novo* prediction of rockfish was carried out with Augustus[14] (the parameter setting as '--uniqueGeneId=true --noInFrameStop=true --gff3=on --genemodel=complete --strand=both'). All evidences of gene model were integrated with Glean[15]. Finally, we identified 24,094 protein-coding genes in rockfish genome (**Table S6**). The average transcript length, CDS length, exon length, exon number and intron length were consistent with homologous species (**Table S7**, **Fig. S4**).

Proteins function of rockfish were obtained from the best Blastp(e-value ≤ 10-5) hit in SwissProt [16] TrEMBL and NCBI NR databases. Gene domain annotation was carried out by searching InterPro database[17]. All genes were aligned against KEGG, and the pathway in which a gene might be involved was identified from the best hits in KEGG[18]. Gene Ontology (GO)[19] terms for genes were obtained from the corresponding InterPro entry (**Table S8, Fig. S5**).

**Genome assembly and gene set assessment**

The short insert library reads were used to map with the assembled genome using BWA software[20] (http://bio-bwa.sourceforge.net/) to statistics the mapping ratio and assess the assembly integrity. 98.13% short reads were mapped to the genome with a coverage of 99.44% explaining the reliability of the genome (**Table S9**).

BUSCO[21] (Benchmarking Universal Single-Copy Orthologs: http://busco.ezlab.org/ ) provides quantitative measures for the assessment of genome assembly completeness, based on evolutionarily-informed expectations of gene content from near-universal single-copy orthologs .There are 93.9% and 94.4% of the 2,586 single copy genes set was found in black rockfish genome and gene set (**Table S10**), indicating the integrity of the genome and gene set.

To check if there is exogenous pollution in rockfish, we conduct a statistical procedure by an in-house program and drew the GC percentage and sequencing depth figure. As shown in the figure, the point is concentrated and no obvious separation which indicating there is no exogenous pollution in the sequencing sample (**Fig. S6**).

**ATAC-seq data analyzing**

Adaptor sequences in ATAC-seq raw data were removed using Cutadapt (https://cutadapt.readthedocs.io/en/stable/index.html) and reads aligned onto rockfish genome using Bowtie2[22] (parameter: -X2000 --local). Reads with the mapping quality less than 30, and reads mapped to the mitochondria genome were filtered out. PCR duplicate reads were removed using Picard's MarkDuplicates (http://broadinstitute.github.io/picard/). Then the model-based analysis of ChIP-seq (MACS version 2.1) was used to call enrichment regions (peaks), with the following settings: callpeak -g 8.11e8 --qvalue 0.05 -p 0.01 --nomodel --shift -75 --extsize 150 -B --SPMR --keep-dup all --call-summits. Peak signal can be visualized in IGV by the Broad Institute (http://software.broadinstitute.org/software/igv/). Peak enrichment and heatmap (**Fig. S7**) were enriched and shown by plotHeatmap provided in deepTools (https://deeptools.readthedocs.io/en/develop/content/tools/plotHeatmap.html)

**RNA sequencing and quantify**

In total, 58 biological RNA sample libraries from 21 tissues were constructed in this study, including heart, liver, spleen, kidney, brain, gill, muscle, intestine, ovarian wall, oocyte in female and heart, liver, spleen, kidney, brain, gill, muscle, intestine, testis, pituitary, genitalia in male randomly selected from twelve healthy 3-year-old fish (six males and six females) (**Table S11**). The transcriptome of embryonic development including 29 RNA-seq libraries belonged to 7 stages (premating, post-mating, pre-fertilization, 1 cell , 32 cell, blastula and gastrulae stage) (**Fig. 1b**, **Table S13** ). RNA libraries were sequenced 100 bp at each end using BGI-seq 500 platform. After removing low-quality reads, the clean reads were quantified using Salmon[23] with default parameters. In order to investigate the regulation network, we used the gene co-expression network constructed by the method of Weighted Gene Co-Expression Network Analysis (WGCNA)[24]. WGCNA (Weighted Gene Co-Expression Network Analysis) is also known as weighted gene co-expression network analysis when dealing with gene expression data[25]. Many functions of WGCNA can also be used for general association networks specified by a symmetric adjacency matrix. Constructing a weighted gene network entails the choice of the soft thresholding power to which co-expression similarity is raised to calculate adjacency. Genes in rockfish were clustered into 28 modules according to gene expression patterns in 21 different tissues (for most tissues, 3 replicates were collected) (**Fig. 2a**), The function of 101 genes which show high correlation with ovarian wall in TM07 is listed in the table **(Table S12)**. Meanwhile, Genes in rockfish were clustered into 39 modules according to gene expression patterns in 7 embryonic development stage (at least 3 replicates were sampled in each time point) (**Fig. 2b**). The co-expression module with correlation value greater than 0.7 and P-value less than 0.05 was selected. After that, 2 modules selected in pre-fertilization and gastrula stage respectively, 1 module selected in post-mating, 32-cell and blastula stage respectively, 0 module selected in the pre-mating ovary and 1-cell embryo**.** Furthermore, we identified a module (3,128 genes in SM20) with a large number of genes co-expressed by the ovary before fertilization (pre-mating, post-mating, and pre-fertilization). The highly expression genes in SM20 were collected and performed an enrichment analysis (**Table S14**).

**Phylogenetic tree construction and divergence time estimate**

The gene family analysis was conducted by Treefam[26] pipeline (version 0.50) using the following steps:

1) Protein sequences (*Danio rerio*, *Gasterosteus aculeatus*, *Hippocampus comes*, *Oreochromis niloticus*, *Oryzias latipes*, *Takifugu rubripes*, *Xiphophorus maculatus Poecilia formosa* , *Cynoglossus semilaevis*, *Astyanax mexicanus*, *Larimichthys crocea*, *Tetraodon nigroviridis*, *Gadus morhua*, *Ietalurus Punetaus*, *Lepisosteus oculatus* and *Callorhinchus milii*) were downloaded from the Ensembl database (Release 92) or NCBI. BLASTP was employed to identify potential homologous genes using E-value< 1e-10.

2) The raw Blast results were refined using solar (an in-house software, version0.9.6) by which the high-scoring segment pairs (HSPs) were conjoined.

3) Similarity between protein sequences were evaluated using bit-score, followedby clustering protein sequences into gene families using hcluster_sg, a hierarchical clustering algorithm in the Treefam pipeline with the parameters "-w 5 -s 0.33 -m 100000".The gene family cluster result of ten species is shown in **Fig. S8.**

Evolutionary analyses were performed using single-copy protein-coding genes from the 16 species. Amino acid and nucleotide sequences of ortholog genes were aligned by the multiple alignment software MUSCLE[27] with the default parameter. A total number of 1761 single-copy ortholog alignments were concatenated into a super alignment matrix of 1,349,014 amino acid. A Maximum Likelihood method (ML) tree was inferred based on the matrix of nucleotide sequences using RAxML[28], applying default nucleotide substitution model-PROTGAMMAAUTO. Clade support was assessed using bootstrapping algorithm in RAxML package with 1,00 alignment replicates (**Fig. S9**).

**Zona pellucida sperm-binding protein gene family**

ZP(zona pellucida) gene sequences of the zebrafish (*Danio rerio*), medaka (*Oryzias latipes*), tilapia (*Oreochromis niloticus*), Amazon molly (*Poecilia Formosa*), platyfish (*Xiphophorus maculates*), tetraodon (*Tetraodon nigroviridis*), stickleback (*Gasterosteus aculeatus*) were collected from Ensemble database according to previous studies[29]. Using the similar manual menthod: ZP-like sequences in black rockfish were identified by blastp (Identity>25, E-value<1e$^{-5}$) against peptide sequences, using ZP sequences of zebrafish and medeka as queries. To reduce redundant matches, two method were used in this research: firstly, the candidate sequence in black rockfish were further used to back search against the NCBI by blastn, then a phylogenetic tree was conducted using candidate sequences and identified reference sequences. After the rigorous filter, 22 zona pellucida sperm-binding genes were left (**Table S15**). Finally, a ML (Maximum Likelihood) tree was conducted with 8 bony fishes ZP genes (**Table S16, Fig S10**. According to the phylogeny tree cluster, ZP genes can be divided into four subfamilies cluster, including ZPB, ZPC, ZPD, ZPAX. The expression quantity of the ZP genes in different tissues and different embryonic development stage was normalized and showed by Heatmap package (**Table S17, Fig. 3b, Fig. S11**).

**Astacin metalloproteinase genes in black rockfish**

The protein sequences of the black rockfish astacin family that were predicted in this study were extracted and manually curated. The genomic protein sequences of 7 bony fishes were downloaded from Ensembl database. The astacin family of zebrafish, medaka, stickleback, tilapia, tetraodon, platyfish were obtained basing on previous study[30] and verified manually. To obtain astacin family genes in rockfish and amazon molly, a verified method was performed: Fistly, a domain prediction pipline was performed to pick the protein sequences which have the astacin protein domain; secondly, the longest transcript was chosen and the short ones which have the same gene ID were removed to avoid duplication; lastly, the candidate protein sequences of astacin family were further filter by NR database using Blastp. After that, 26 astacin family genes in rockfish and 11 in amazon molly were left **(Table S18,Table S19)**. The phylogeny tree was constructed using ML(Maximum Likelihood) method imply in Fasttree[31] (**Fig. 4a**). Astacin family genes can be devided into five subfamilies cluster, including Patristacin/astacin, Nephrosin, Choriolysin, HCE1-like and HCE2-like according to the phylogeny tree. Phylogenetic analysis found two subfamilies (Hatching enzyme and HCE1-like) trend to expand in rockfish, suggesting that the expansion is specific to rockfish. The expression pattern of the astacin genes in different tissues and different embryonic development stage was analyzed by Heatmap in R (version 3.4.1).

#### Supplementary Figures

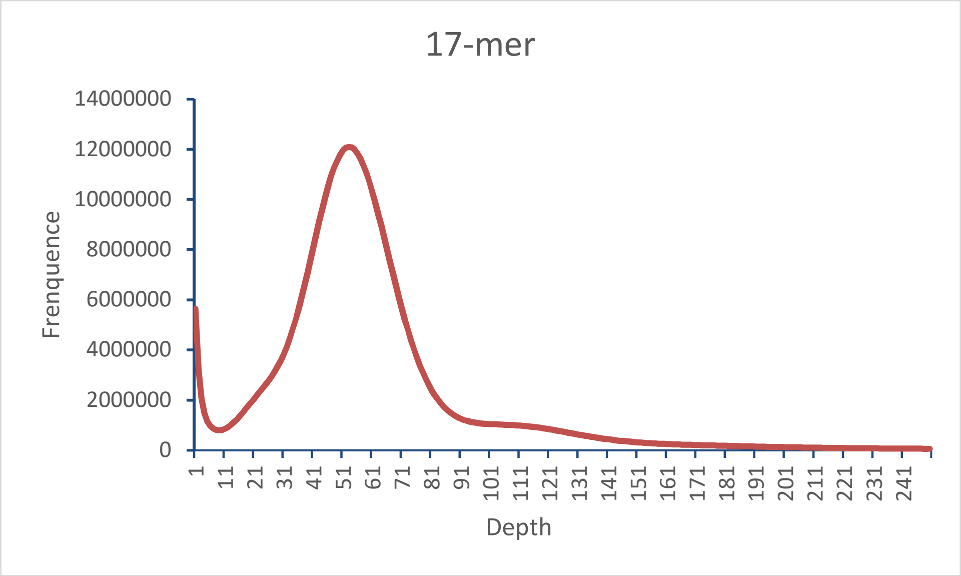


**Fig. S1.** Genome size and depth estimate based on K-mer spectrum. The peak depth is 57 and the total K-mer number is 49,484,519,640.

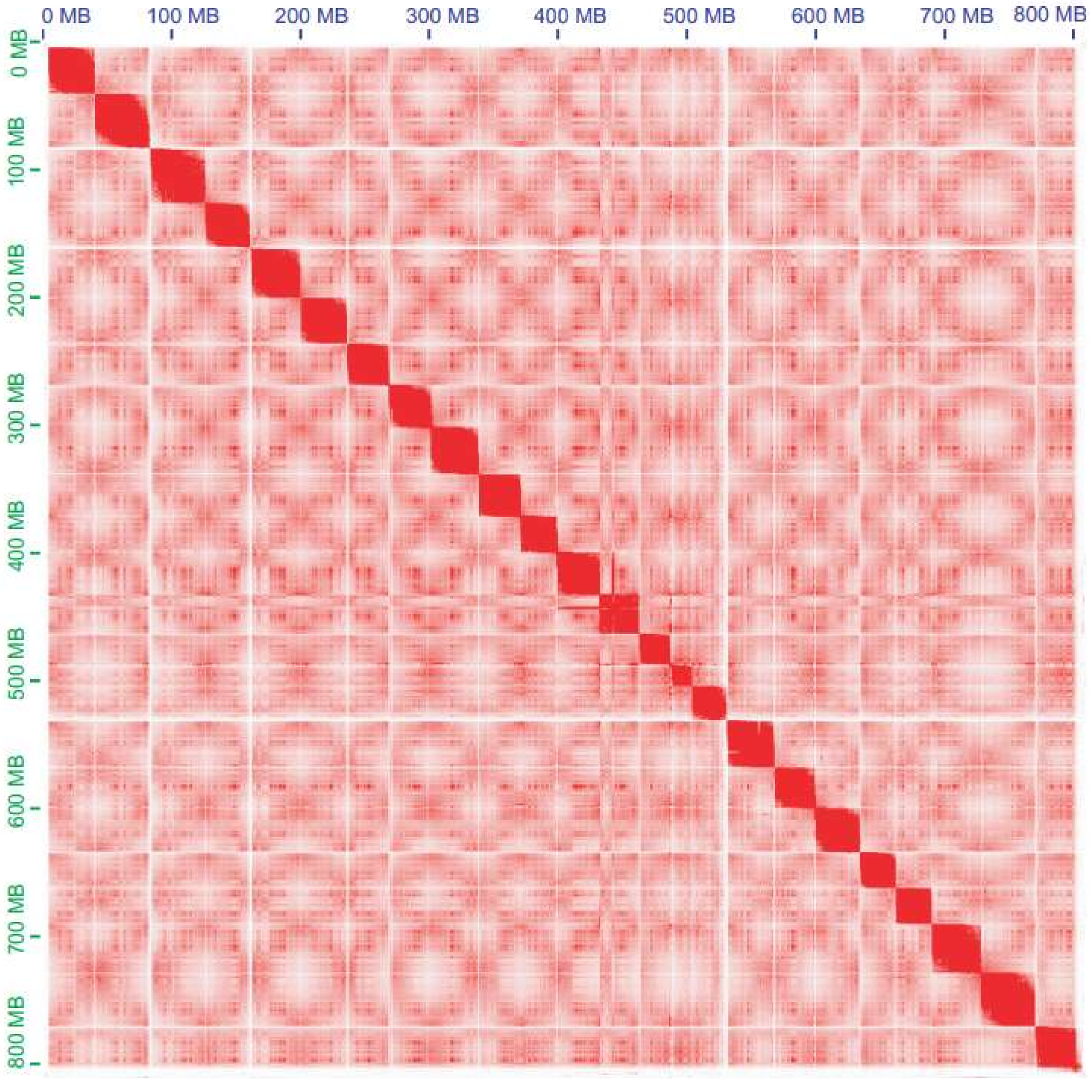


**Fig. S2.** Heatmap of chromosome interaction intensity in Hi-C assembly. The X axis and Y axis both are the length of rockfish chromosome.

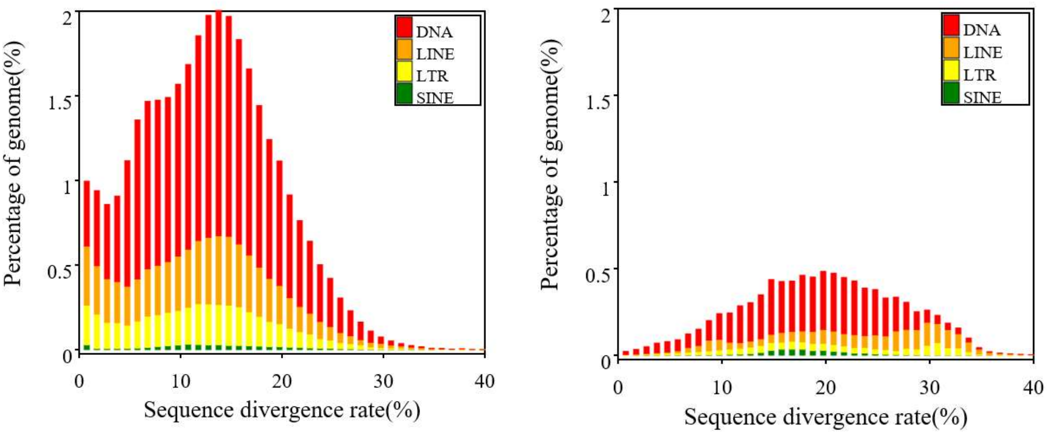


De novo Homolog

**Fig. S3.** Distribution of TEs in the rockfish genome. The TEs included DNA transposons (DNA) and RNA transposons (including LINE, LTR, SINE).

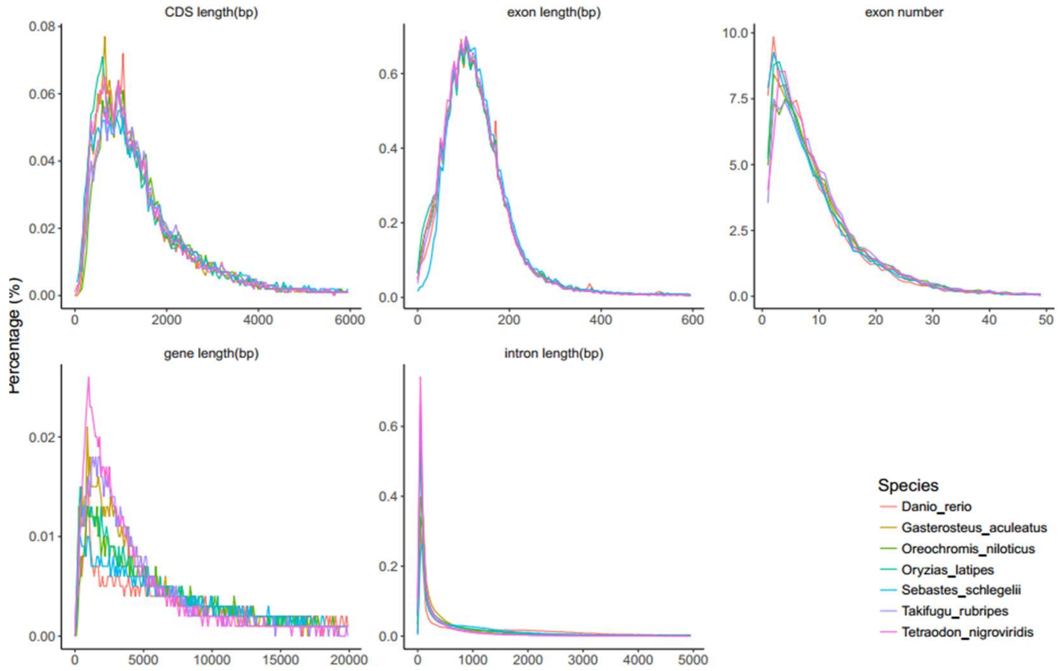


**Fig. S4.** Comparison of gene structure among homolog species. The length of mRNA, exon, intron and CDS and the number of exon among homolog species were shown respectively. The homolog species selected were shown with the following abbreviation with different color.

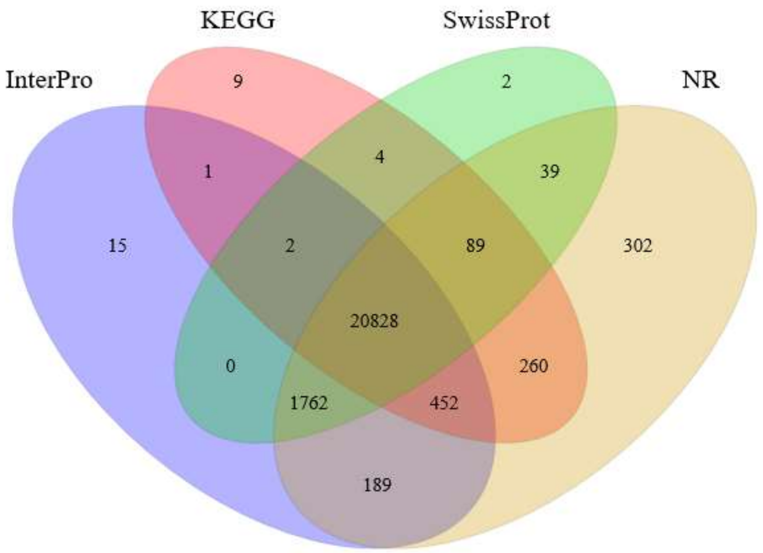


**Fig. S5.** Comparison of the gene sets annotated by different database (InterPro, KEGG, SwissProt, NR).

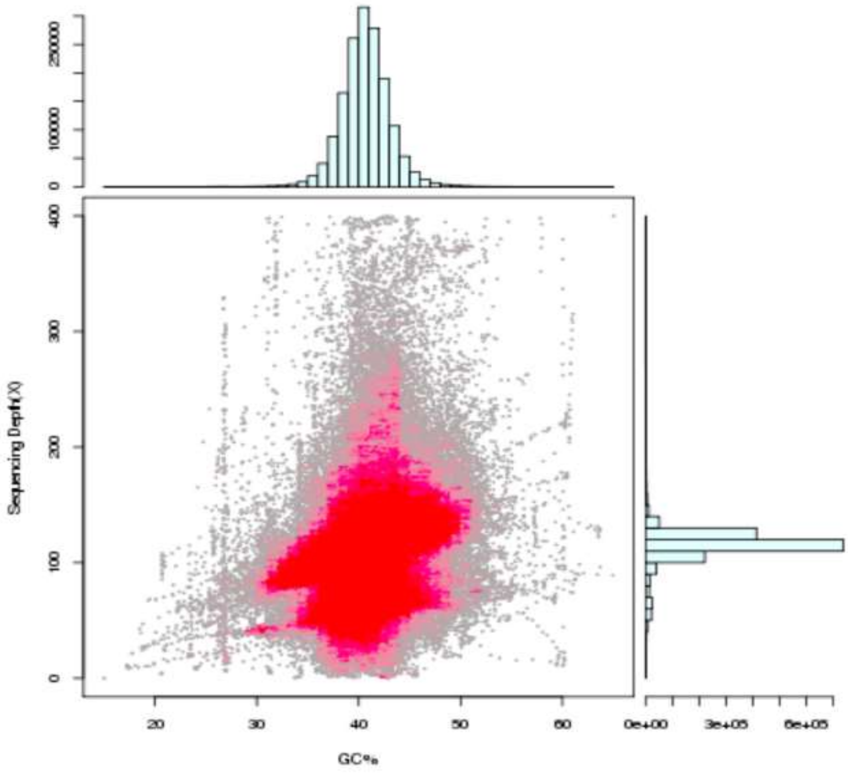


**Fig. S6.** The GC percentage and sequencing depth figure. No obvious GC separation indicating the sample is clean.

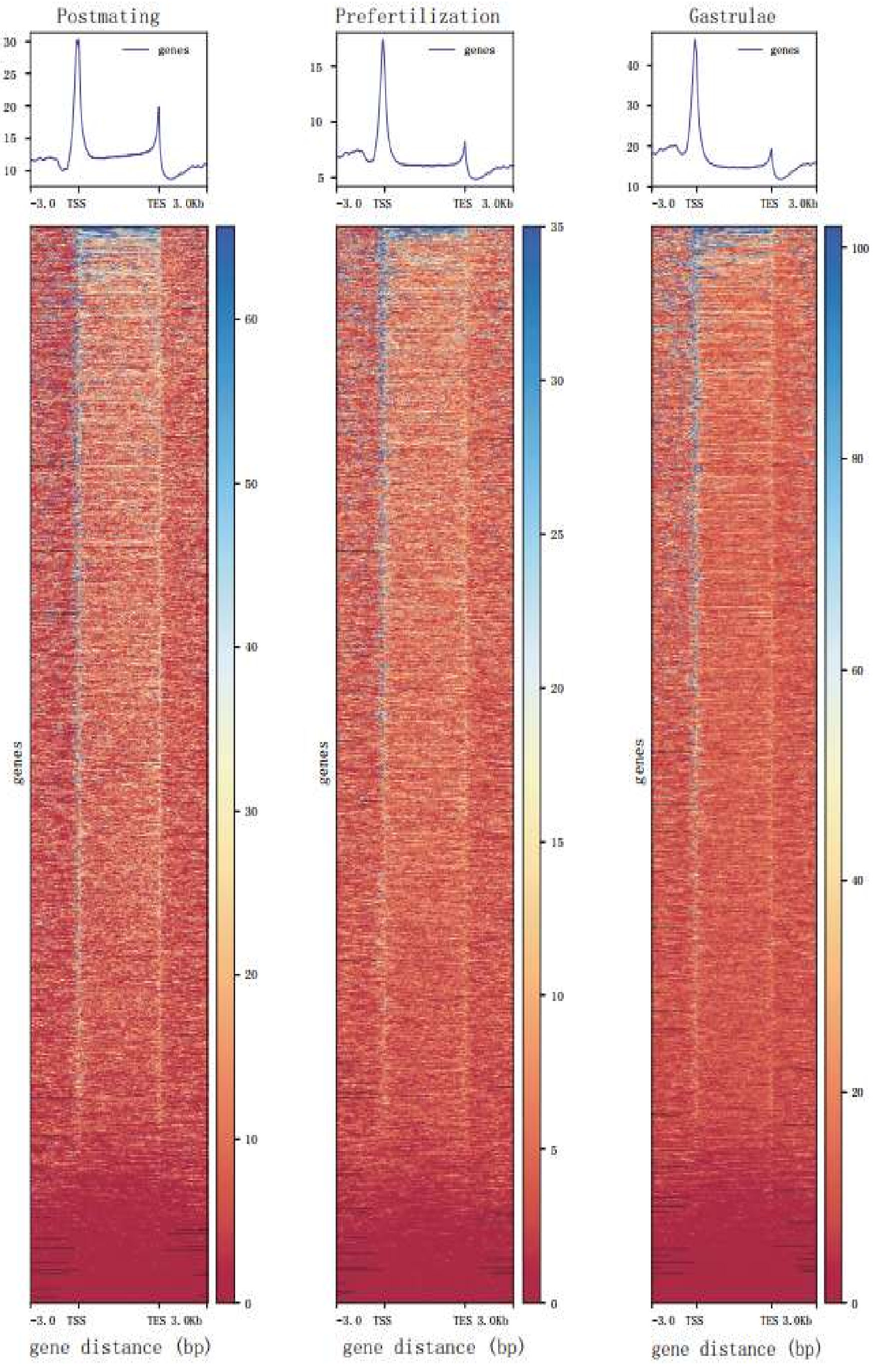


**Fig. S7.** The average ATAC-seq enrichment and heatmap at three stage (postmating, prefertilization and gastrulae)

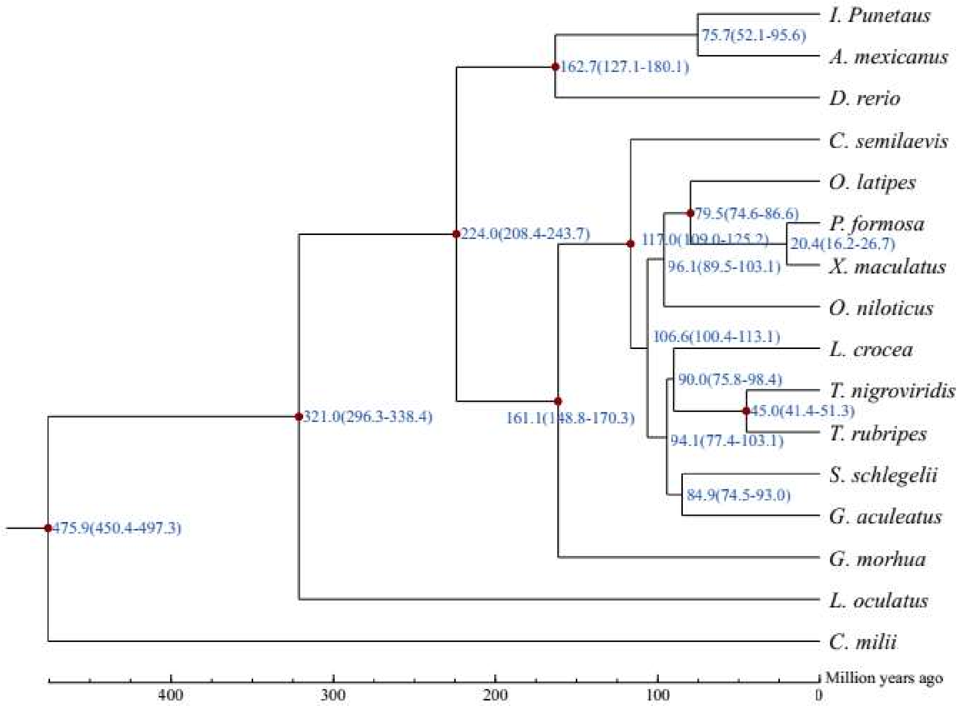


**Fig. S9.** The estimation of divergence time. The numbers on the nodes (blue) represented the divergence times from present (million years ago, Mya) and the numbers in parentheses represented the range of divergence times. The red dot was the time correct point we chose from Timetree or fossil calibrations.

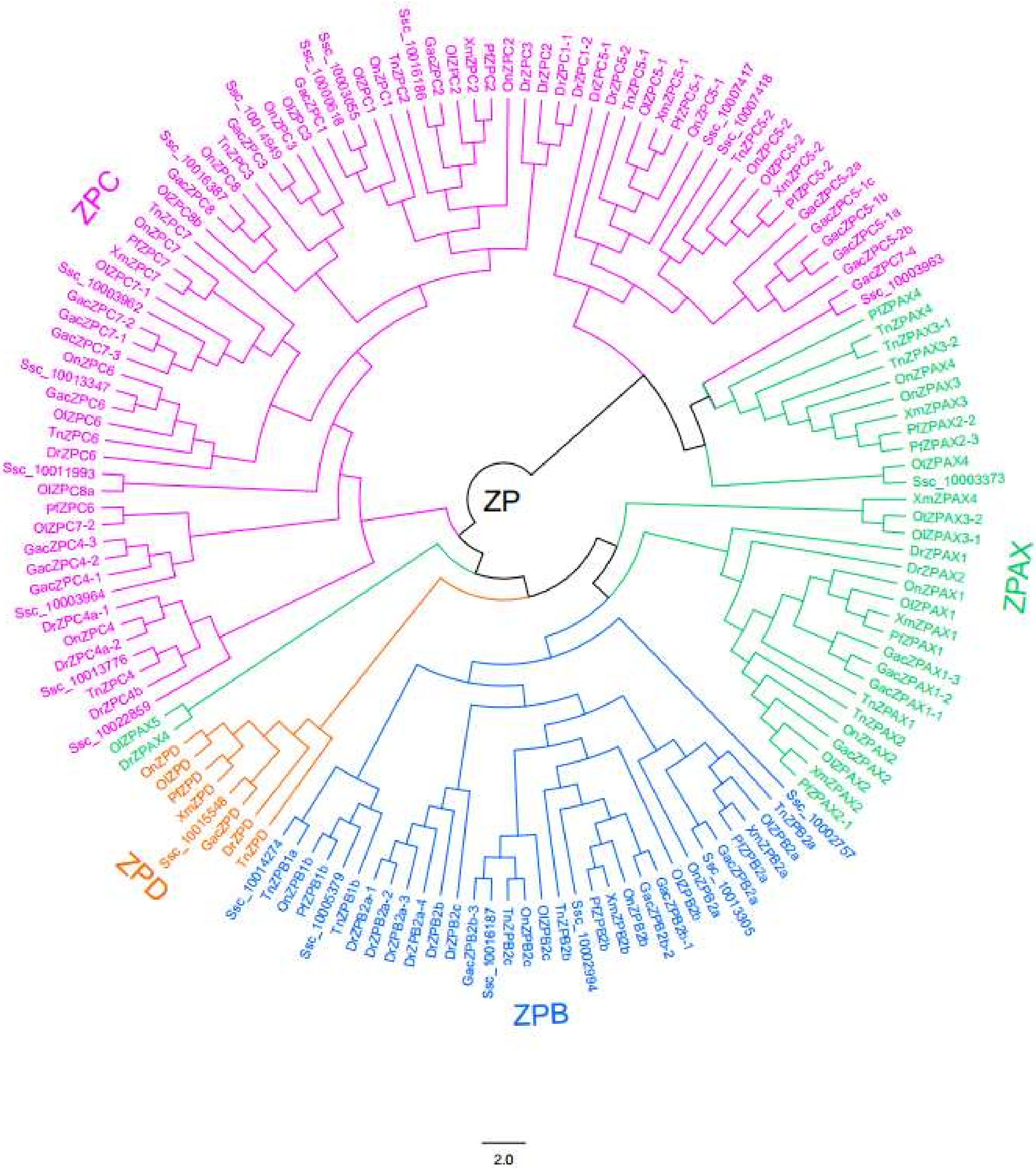


**Fig. S10.** Phylogenetic tree constructed with protein sequences encoded by various ZP genes in 8 bony fish

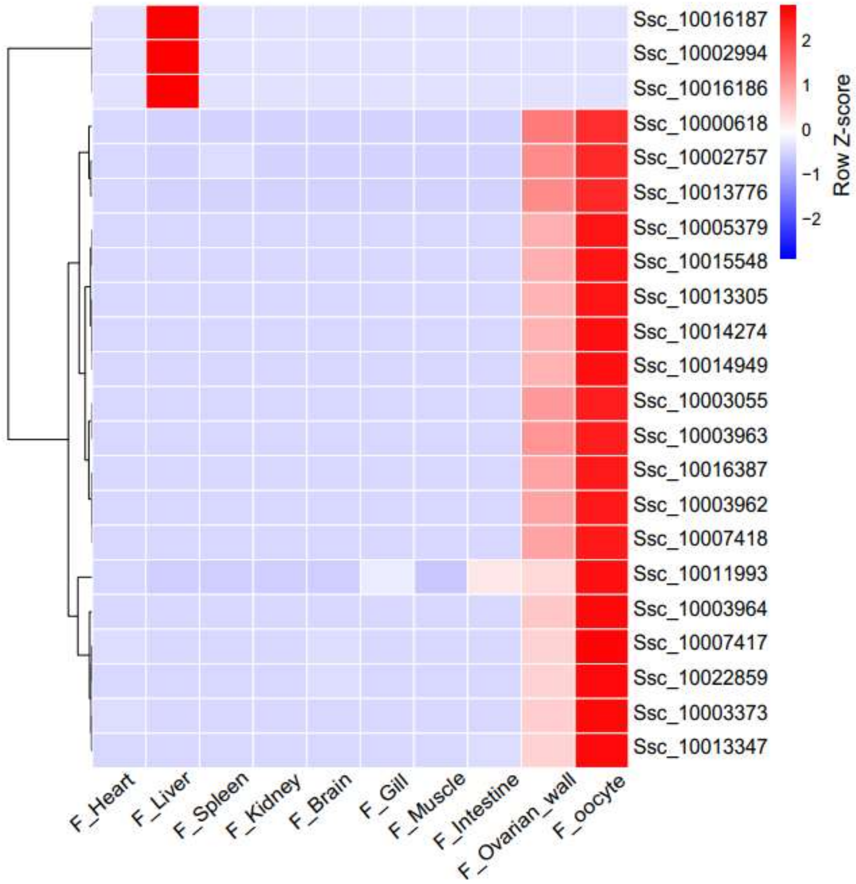


**Fig. S11.** The expression profile of ZP genes in black rockfish different tissues

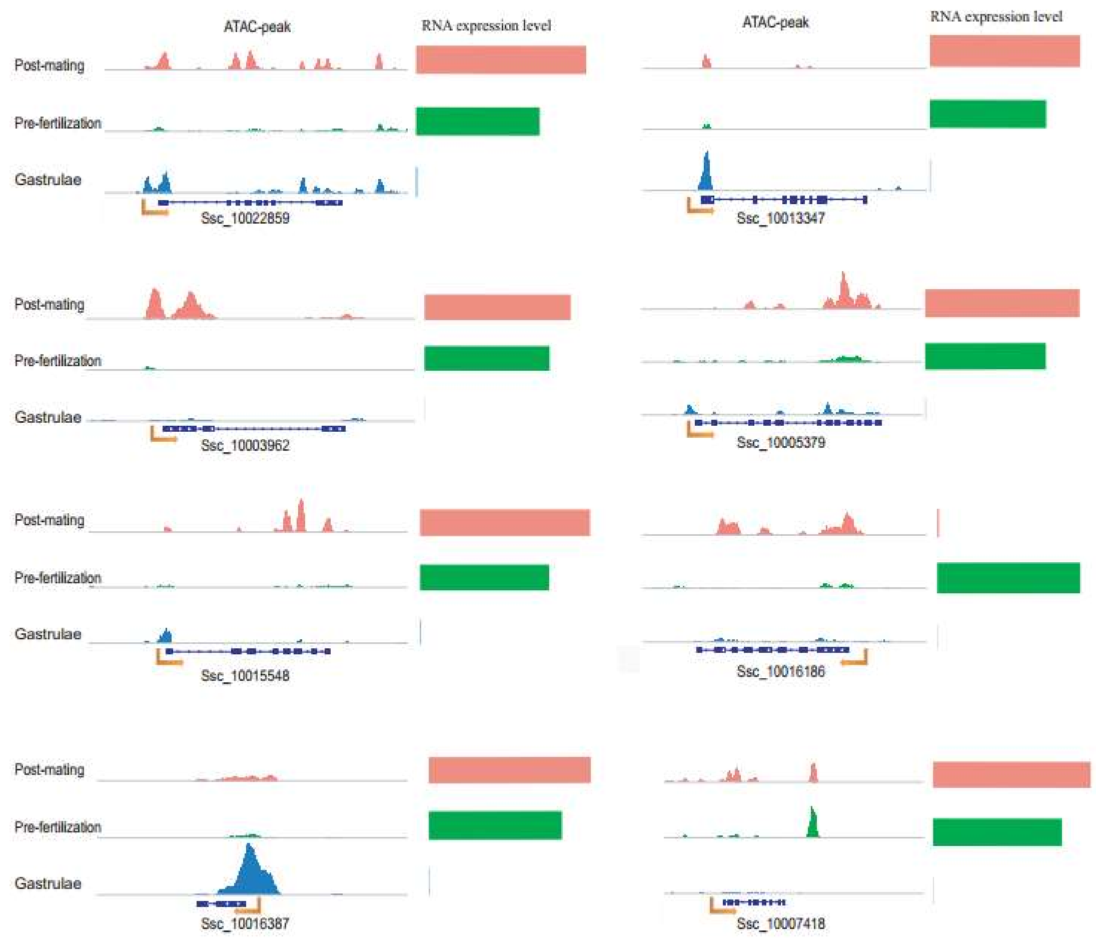


**Fig. S12** Signals of accessible chromatin and associated RNA expression levels of selected ZP genes. ATAC result of each gene is shown on the left, with peaks indicating accessible chromatin regions; gene expression levels (in TPM) on the right.

### Supplementary Tables

**Table S1.** The table of genome size and depth estimate

| K | K-mer_num | Peak_depth | Genome Size | Used Bases | Used Reads | Depth |
|---|---|---|---|---|---|---|
| 17 | 49,484,519,640 | 57 | 868,149,467 | 55,393,119,000 | 369,287,460 | 63.81 |

**Table S2.** Statistics of the genome sequencing data of rockfish

| | **Average Read Length/bp** | **BaseNum/Gbp** | **Sequence Depth*** |
|---|---|---|---|
| BGI-seq500 | 150 | 98.1 | 114.6 |
| PacBio | 11,874.2 | 57.3 | 66.0 |
| HI-C | 100 | 152.9 | 188.8 |
| Total | - | 308.3 | 355.2 |

*Based on clean data and the estimated genome size of 868Mb

**Table S3.** Statistics of rockfish genome assembly.

| | Contig | | Scaffold | |
|---|---|---|---|---|
| | **Size(bp)** | **Number** | **Size(bp)** | **Number** |
| N90 | 288,836 | 298 | 288,836 | 296 |
| N80 | 1,189,441 | 167 | 1,189,441 | 165 |
| N70 | 2,011,124 | 117 | 2,058,930 | 115 |
| N60 | 3,036,147 | 85 | 3,048,075 | 83 |
| N50 | 3,845,851 | 60 | 3,847,635 | 60 |
| Longest | 16,708,100 | ---- | 16,708,100 | ---- |
| Total* | 811,127,112 | 1,454 | 811,127,114 | 1,452 |

*Only contigs with lengths >= 2000 bp were included in the genome assembly;

**Table S4.** The pseudochromosomes length after Hi-C assembly. The pseudochromosomes are ordered from long to short in length.

| Pseudochromosomes | length(bp) | percentage |
|---|---|---|
| chr_1 | 43,928,042 | 5.40% |
| chr_2 | 42,604,752 | 5.24% |
| chr_3 | 41,099,864 | 5.06% |
| chr_4 | 40,690,939 | 5.01% |
| chr_5 | 39,638,966 | 4.88% |
| chr_6 | 38,184,723 | 4.70% |
| chr_7 | 37,532,197 | 4.62% |
| chr_8 | 36,675,045 | 4.51% |
| chr_9 | 36,546,961 | 4.50% |
| chr_10 | 36,163,463 | 4.45% |
| chr_11 | 34,771,888 | 4.28% |
| chr_12 | 33,217,404 | 4.09% |
| chr_13 | 32,715,586 | 4.02% |
| chr_14 | 32,366,827 | 3.98% |
| chr_15 | 31,180,171 | 3.84% |
| chr_16 | 30,925,316 | 3.80% |
| chr_17 | 30,896,493 | 3.80% |
| chr_18 | 30,831,959 | 3.79% |
| chr_19 | 30,050,174 | 3.70% |
| chr_20 | 29,890,757 | 3.68% |
| chr_21 | 28,468,378 | 3.50% |
| chr_22 | 28,125,667 | 3.46% |
| chr_23 | 27,621,098 | 3.40% |
| chr_24 | 16,577,481 | 2.04% |
| unmap | 1,115,963 | 0.14% |
| Total | 812936077 | 100% |

**Table S5.** The prediction of repeats elements in rockfish genome.

| | Repbase TEs | | TE protiens | | De novo | | Combined TEs | |
|---|---|---|---|---|---|---|---|---|
| Type | Length (bp) | % in genome | Length (bp) | % in genome | Length (bp) | % in genome | Length (bp) | % in genome |
| DNA | 37,322,080 | 4.60 | 2,441,317 | 0.30 | 179,554,256 | 22.14 | 194,009,383 | 23.92 |
| LINE | 15,488,614 | 1.91 | 9,444,767 | 1.16 | 68,579,110 | 8.45 | 77,585,678 | 9.57 |
| SINE | 2,461,206 | 0.30 | 0 | 0.00 | 4,587,535 | 0.57 | 6,746,377 | 0.83 |
| LTR | 7,921,660 | 0.98 | 2,634,056 | 0.32 | 46,097,593 | 5.68 | 51,133,018 | 6.30 |
| Other | 17,640 | 0.00 | 0 | 0.00 | 0 | 0.00 | 17,640 | 0.00 |
| Unknown | 0 | 0.00 | 0 | 0.00 | 12,159,800 | 1.50 | 12,159,800 | 1.50 |
| Total | 57217352 | 7.05 | 14,516,194 | 1.79 | 274,309,843 | 33.82 | 287,072,331 | 35.39 |

**Table S6.** Summary of predicted gene models in the rockfish genome. The final gene set of rockfish genome was created by merging AUGUST prediction, homologous predictions and RNA-seq data.

| #Gene set | Number of genes | CDS+intron len | CDS len | exon len | intron len | Exons per gene |
|---|---|---|---|---|---|---|
| *Danio_rerio* | 22,362 | 26,760.44 | 1,587.11 | 180.07 | 3,221.6 | 8.81 |
| *Takifugu_rubripes* | 20,164 | 15,362.7 | 1,620.55 | 168.04 | 1,589.84 | 9.64 |
| *Tetraodon_nigroviridis* | 20,833 | 13,291.67 | 1,471.92 | 162.84 | 1,470.28 | 9.04 |
| *Gasterosteus_aculeatus* | 22,473 | 13,440.13 | 1,508.84 | 167.03 | 1,485.19 | 9.03 |
| *Larimichthys_crocea* | 23,723 | 15,797.15 | 1,674.15 | 173.28 | 1,630.49 | 9.66 |
| *Cynoglossus semilaevis* | 21,478 | 16,855.58 | 1,717.88 | 173.53 | 1,700.97 | 9.9 |
| *Oreochromis niloticus* | 22,793 | 17,097.07 | 1,665.2 | 173.79 | 1,798.24 | 9.58 |
| *Oryzias latipes* | 20,554 | 13,921.36 | 1,503.46 | 168.74 | 1,569.95 | 8.91 |
| *Poecilia formosa* | 24,057 | 16,035.14 | 1,639.19 | 177.00 | 1,742.69 | 9.26 |
| AUGUST | 18,255 | 14,514.94 | 1,456.16 | 175.03 | 1,784.11 | 8.32 |
| GLEAN* | 24,094 | 15,781.76 | 1,709.42 | 175.52 | 1,610.3 | 9.74 |

* The GLEAN gene set contains the integrated result of de novo genes predictions, homolog-based genes predictions and transcriptome-based annotation by using GLEAN software.

**Table7.** Genetic structure characteristics in rockfish **c**omparing with homolog species

| Gene set | Number of genes | CDS+intron len(avg) | CDS len(avg) | exon len(avg) | intron len(avg) | Exons per gene(avg) |
|---|---|---|---|---|---|---|
| *Sebastes schlegelii* | 22,011 | 18,368.79 | 1,732.01 | 175.51 | 1,875.91 | 9.87 |
| *Danio rerio* | 25,619 | 25,207.59 | 1,642.64 | 174.39 | 2,798.97 | 9.42 |
| *Oryzias latipes* | 19,699 | 12,145.58 | 1,515.82 | 147.82 | 1,148.61 | 10.25 |
| *Tetraodon nigroviridis* | 19,602 | 6,066.17 | 1,516.59 | 144.2 | 478.02 | 10.52 |
| *Oreochromis niloticus* | 21,437 | 14,903.11 | 1,714.22 | 157.25 | 1,332.07 | 10.9 |
| *Gasterosteus aculeatus* | 20,787 | 8,451.06 | 1,548.67 | 148.94 | 734.44 | 10.4 |
| *Takifugu rubripes* | 18,523 | 7,492.75 | 1,693.53 | 152.61 | 574.33 | 11.1 |

**Table S8.** The number of gene model that can be annotated using different databases.

| | Total | Nr | Swissprot | KEGG | TrEMBL | Interpro | GO | Overall |
|---|---|---|---|---|---|---|---|---|
| Number | 24,094 | 23,921 | 22,726 | 21,645 | 23,985 | 23,249 | 17,551 | 24,025 |
| Percentage | 100% | 99.28% | 94.32% | 89.84% | 99.55% | 96.49% | 72.84% | 99.71% |

**Table S9.** Assessment the genome coverage rate using raw data. Mapping rate was calculated through raw reads mapped to the genome to explain the reliability of the genome coverage.

| | Result |
|---|---|
| Read mapping rate (%) | 98.13% |
| Genome average sequencing depth (X) | 116.08 |
| Coverage of genome (%) | 99.44% |
| Coverage of genome > 4X (%) | 99.18% |
| Coverage of genome > 10X (%) | 98.84% |
| Coverage of genome > 20X (%) | 98.29% |

**Table S10.** BUSCO results of rockfish genome and gene set.

| | Genome BUSCO | Gene BUSCO |
|---|---|---|
| Complete BUSCOs | 93.9% | 94.4% |
| Complete Single-Copy BUSCOs | 91.7% | 91.8% |
| Complete Duplicated BUSCOs | 2.2% | 2.6% |
| Fragmented BUSCOs | 3.0% | 3.1% |
| Missing BUSCOs | 3.1% | 2.5% |

**Table S14.** Enrichment analysis of high expression genes in SM20 using gene ontology, KEGG pathway and InterPro protein domain information. Molecular function and cellular component are two aspects of gene ontology.

| Pathway ID | Pathway description | Count | FDR |
|---|---|---|---|
| | **Molecular Function (GO)** | | |
| GO:0003723 | RNA binding | 30 | 0.00349 |
| | **Cellular Component (GO)** | | |
| GO:0005575 | Cellular component | 230 | 5.86E-07 |
| GO:0005622 | intracellular | 192 | 5.86E-07 |
| GO:0043227 | membrane-bounded organelle | 156 | 7.85E-07 |
| GO:0005623 | cell | 201 | 8.43E-07 |
| GO:0043226 | organelle | 167 | 8.43E-07 |
| GO:0043229 | intracellular organelle | 165 | 8.43E-07 |
| GO:0044422 | organelle part | 99 | 8.43E-07 |
| GO:0044424 | intracellular part | 187 | 8.43E-07 |
| GO:0044446 | intracellular organelle part | 97 | 8.43E-07 |
| GO:0044464 | cell part | 201 | 8.43E-07 |
| GO:0043231 | intracellular membrane-bounded organelle | 152 | 1.12E-06 |
| GO:0044428 | nuclear part | 39 | 4.60E-05 |
| GO:0070013 | intracellular organelle lumen | 35 | 9.17E-05 |
| GO:0031974 | membrane-enclosed lumen | 36 | 9.19E-05 |
| GO:0005737 | cytoplasm | 128 | 0.000103 |
| GO:0031967 | organelle envelope | 27 | 0.000119 |
| GO:0032991 | macromolecular complex | 60 | 0.000125 |
| GO:0005739 | mitochondrion | 33 | 0.000284 |
| GO:0005634 | nucleus | 94 | 0.00108 |
| GO:0031981 | nuclear lumen | 28 | 0.0015 |
| GO:0044429 | mitochondrial part | 23 | 0.00224 |
| GO:0031090 | organelle membrane | 47 | 0.00287 |

| | | | |
|---|---|---|---|
| GO:0016020 | membrane | 89 | 0.00294 |
| GO:0005740 | mitochondrial envelope | 20 | 0.00397 |
| GO:0031966 | mitochondrial membrane | 19 | 0.00499 |
| GO:0043232 | intracellular non-membrane-bounded organelle | 45 | 0.00529 |
| GO:0043234 | protein complex | 45 | 0.00606 |
| GO:0005654 | nucleoplasm | 18 | 0.0112 |
| GO:0005681 | spliceosomal complex | 7 | 0.0154 |
| GO:0030529 | ribonucleoprotein complex | 18 | 0.0177 |
| GO:0019866 | organelle inner membrane | 14 | 0.0215 |
| GO:0044451 | nucleoplasm part | 15 | 0.0253 |
| GO:0044425 | membrane part | 71 | 0.0379 |
| GO:0005743 | mitochondrial inner membrane | 13 | 0.0406 |
| GO:0044444 | cytoplasmic part | 75 | 0.0406 |
| | **KEGG Pathways** | | |
| 190 | Oxidative phosphorylation | 38 | 7.28E-13 |
| 3040 | Spliceosome | 34 | 1.61E-10 |
| 1100 | Metabolic pathways | 131 | 4.59E-07 |
| 3015 | mRNA surveillance pathway | 19 | 0.000132 |
| 970 | Aminoacyl-tRNA biosynthesis | 12 | 0.000161 |
| 3008 | Ribosome biogenesis in eukaryotes | 16 | 0.000587 |
| 3013 | RNA transport | 24 | 0.00102 |
| 3060 | Protein export | 8 | 0.00102 |
| 4145 | Phagosome | 21 | 0.00649 |
| 3050 | Proteasome | 11 | 0.00682 |
| 3022 | Basal transcription factors | 9 | 0.0148 |
| 4142 | Lysosome | 19 | 0.0181 |
| 1200 | Carbon metabolism | 17 | 0.022 |
| 4110 | Cell cycle | 19 | 0.022 |

| | | | |
|---|---|---|---|
| 1120 | Microbial metabolism in diverse environments | 21 | 0.0242 |
| 310 | Lysine degradation | 11 | 0.0302 |
| 5168 | Herpes simplex infection | 21 | 0.0309 |
| 30 | Pentose phosphate pathway | 7 | 0.0319 |
| 4540 | Gap junction | 16 | 0.035 |
| 4512 | ECM-receptor interaction | 12 | 0.0409 |
| 4320 | Dorso-ventral axis formation | 6 | 0.0471 |
| 3460 | Fanconi anemia pathway | 9 | 0.0489 |
| | **InterPro Protein Domains** | | |
| IPR001507 | Zona pellucida domain | 16 | 0.0276 |

**Table S15.** Number variation of Zone pellucid (ZP) genes in 8 bony fish.

| | **ZPB** | **ZPC** | **ZPD** | **ZPAX** | **Total** |
|---|---|---|---|---|---|
| ***Sebastes schlegelii*** | 6 | 14 | 1 | 1 | 22 |
| ***Danio rerio*** | 6 | 10 | 1 | 3 | 20 |
| ***Oryzias latipes*** | 3 | 10 | 1 | 6 | 20 |
| ***Oreochromis niloticus*** | 4 | 9 | 1 | 4 | 18 |
| ***Poecilia formosa*** | 4 | 5 | 1 | 4 | 14 |
| ***Xiphophorus maculatus*** | 2 | 4 | 1 | 4 | 11 |
| ***Tetraodon nigroviridis*** | 5 | 7 | 1 | 5 | 18 |
| ***Gasterosteus aculeatus*** | 4 | 17 | 1 | 4 | 26 |

**Table S17.** TPM(Transcripts Per Million) of 22 ZP genes in seven stages during reproduction

| Gene_ID | Pre-mating | Post-mating | Pre-fertilization | 1 cell | 32 cell | Blastula | Gastrula |
|---|---|---|---|---|---|---|---|
| Ssc_10000618 | 51.22 | 56.87 | 61.92 | 6.31 | 2.27 | 1.41 | 0.39 |
| Ssc_10002757 | 10.81 | 12.33 | 10.89 | 0.96 | 1.14 | 2.19 | 0.06 |
| Ssc_10002994 | 0.13 | 0.31 | 1.26 | 0.06 | 0.07 | 0.00 | 0.07 |
| Ssc_10003055 | 108.70 | 125.79 | 106.01 | 10.42 | 6.09 | 6.29 | 0.56 |
| Ssc_10003373 | 20.62 | 19.01 | 14.29 | 0.82 | 0.31 | 0.26 | 0.11 |
| Ssc_10003962 | 617.67 | 639.66 | 546.06 | 47.02 | 3.70 | 6.86 | 1.09 |
| Ssc_10003963 | 1874.10 | 1685.22 | 1418.86 | 116.93 | 11.50 | 61.55 | 2.93 |
| Ssc_10003964 | 470.45 | 480.46 | 343.80 | 36.68 | 11.69 | 19.69 | 1.21 |
| Ssc_10005379 | 320.96 | 322.92 | 253.95 | 17.73 | 5.35 | 9.70 | 1.10 |
| Ssc_10007417 | 142.05 | 121.27 | 90.52 | 5.59 | 2.74 | 6.25 | 0.19 |
| Ssc_10007418 | 1869.43 | 2134.98 | 1748.59 | 121.44 | 34.70 | 72.94 | 6.67 |
| Ssc_10011993 | 30.90 | 37.26 | 28.20 | 4.76 | 6.33 | 9.93 | 0.55 |
| Ssc_10013305 | 2375.50 | 2380.98 | 1609.35 | 124.28 | 23.83 | 80.66 | 2.88 |
| Ssc_10013347 | 1055.51 | 892.48 | 692.71 | 77.73 | 162.27 | 224.25 | 3.90 |
| Ssc_10013776 | 2759.21 | 2700.88 | 2370.41 | 212.80 | 97.33 | 116.94 | 10.34 |
| Ssc_10014274 | 2710.03 | 2596.74 | 2070.27 | 158.96 | 123.43 | 171.66 | 12.18 |
| Ssc_10014949 | 471.68 | 469.47 | 352.30 | 26.10 | 9.95 | 26.54 | 1.54 |
| Ssc_10015548 | 309.22 | 358.48 | 271.81 | 27.12 | 12.93 | 15.21 | 1.97 |
| Ssc_10016186 | 0.01 | 0.03 | 2.32 | 0.00 | 0.00 | 0.00 | 0.00 |
| Ssc_10016187 | 0.66 | 0.47 | 10.61 | 0.00 | 0.05 | 0.10 | 0.01 |
| Ssc_10016387 | 165.95 | 220.11 | 180.47 | 19.06 | 8.58 | 46.59 | 0.83 |
| Ssc_10022859 | 167.27 | 152.54 | 110.75 | 13.54 | 7.62 | 10.71 | 1.26 |

**Table S18.** Number variation of astacin family genes in 8 bony fish

| | Viviparity | PA** | HE | H1L | H2L | Nep | Total |
|---|---|---|---|---|---|---|---|
| Rockfish* | Y | 1 | 8 | 12 | 2 | 3 | 26 |
| Molly | Y | 7 | 0 | 1 | 2 | 1 | 11 |
| Platyfish | Y | 6 | 0 | 2 | 2 | 1 | 11 |
| Medaka | N | 0 | 2 | 5 | 2 | 1 | 10 |
| Zebrafish | N | 0 | 3 | 2 | 2 | 2 | 9 |
| Stickleback | N | 0 | 5 | 4 | 1 | 2 | 12 |
| Tilapia | N | 2 | 3 | 3 | 2 | 1 | 11 |
| Tetraodon | N | 2 | 1 | 3 | 2 | 1 | 9 |

*Rockfish: *Sebastes schlegelii;* Molly: *Poecilia formosa*; Platyfish: *Xiphophorus maculatus*; Medaka: *Oryzias latipes*; Zebrafish: *Danio rerio*; Stickleback: *Gasterosteus aculeatus*; Tilapia: *Oreochromis niloticus*; Tetraodon: *Tetraodon nigroviridis.*

**PA: Patristacin/Astacin; HE: Hatching enzyme; H1L: HCE1-like; H2L: HCE2-like; Nep: Nephrosin;

**Table S19.** Gene IDs of astacin family genes in 8 teleost genomes

| Species | Astacin_Gene_ID | Species | Astacin_Gene_ID |
|---|---|---|---|
| Zebrafish | ENSDARG00000010423 | Platyfish | ENSXMAG00000000793 |
| Zebrafish | ENSDARG00000019122 | Platyfish | ENSXMAG00000000804 |
| Zebrafish | ENSDARG00000022670 | Platyfish | ENSXMAG00000015480 |
| Zebrafish | ENSDARG00000023656 | Platyfish | ENSXMAG00000015483 |
| Zebrafish | ENSDARG00000024503 | Platyfish | ENSXMAG00000017307 |
| Zebrafish | ENSDARG00000052578 | Platyfish | ENSXMAG00000017982 |
| Zebrafish | ENSDARG00000058409 | Platyfish | ENSXMAG00000017989 |
| Zebrafish | ENSDARG00000069216 | Platyfish | ENSXMAG00000017991 |
| Zebrafish | ENSDARG00000070011 | Platyfish | ENSXMAG00000017994 |
| Stickleback | ENSGACG00000000109 | Platyfish | ENSXMAG00000018023 |
| Stickleback | ENSGACG00000000834 | Platyfish | ENSXMAG00000018029 |
| Stickleback | ENSGACG00000003057 | Tetraodon | ENSTNIG00000000310 |
| Stickleback | ENSGACG00000005978 | Tetraodon | ENSTNIG00000004524 |
| Stickleback | ENSGACG00000008997 | Tetraodon | ENSTNIG00000006100 |
| Stickleback | ENSGACG00000012854 | Tetraodon | ENSTNIG00000006101 |
| Stickleback | ENSGACG00000014406 | Tetraodon | ENSTNIG00000010978 |
| Stickleback | ENSGACG00000014411 | Tetraodon | ENSTNIG00000016065 |
| Stickleback | ENSGACG00000014415 | Tetraodon | ENSTNIG00000016473 |
| Stickleback | ENSGACG00000014420 | Tetraodon | ENSTNIG00000016474 |
| Stickleback | ENSGACG00000015307 | Tetraodon | ENSTNIG00000018786 |
| Stickleback | ENSGACG00000015309 | Tetraodon | ENSTNIG00000018787 |
| Medaka | ENSORLG00000000220 | Rockfish | Ssc_10001812 |
| Medaka | ENSORLG00000000239 | Rockfish | Ssc_10001813 |
| Medaka | ENSORLG00000014846 | Rockfish | Ssc_10001814 |
| Medaka | ENSORLG00000014863 | Rockfish | Ssc_10001815 |
| Medaka | ENSORLG00000014873 | Rockfish | Ssc_10001816 |
| Medaka | ENSORLG00000015017 | Rockfish | Ssc_10001817 |
| Medaka | ENSORLG00000016557 | Rockfish | Ssc_10001818 |
| Medaka | ENSORLG00000016562 | Rockfish | Ssc_10001819 |
| Medaka | ENSORLG00000019231 | Rockfish | Ssc_10005763 |
| Medaka | ENSORLG00000019499 | Rockfish | Ssc_10005764 |
| Nile Tilapia | ENSONIG00000000840 | Rockfish | Ssc_10005765 |
| Nile Tilapia | ENSONIG00000003919 | Rockfish | Ssc_10005766 |
| Nile Tilapia | ENSONIG00000003921 | Rockfish | Ssc_10008383 |
| Nile Tilapia | ENSONIG00000006941 | Rockfish | Ssc_10008384 |
| Nile Tilapia | ENSONIG00000009020 | Rockfish | Ssc_10011519 |
| Nile Tilapia | ENSONIG00000009023 | Rockfish | Ssc_10012428 |
| Nile Tilapia | ENSONIG00000015038 | Rockfish | Ssc_10012458 |
| Nile Tilapia | ENSONIG00000016386 | Rockfish | Ssc_10018443 |
| Nile Tilapia | ENSONIG00000018108 | Rockfish | Ssc_10018444 |
| Nile Tilapia | ENSONIG00000020919 | Rockfish | Ssc_10018478 |
| Nile Tilapia | ENSONIG00000020921 | Rockfish | Ssc_10018480 |

| | | | |
|---|---|---|---|
| Amazon molly | ENSPFOG00000003733 | Rockfish | Ssc_10018494 |
| Amazon molly | ENSPFOG00000003954 | Rockfish | Ssc_10019406 |
| Amazon molly | ENSPFOG00000007022 | Rockfish | Ssc_10019957 |
| Amazon molly | ENSPFOG00000007073 | Rockfish | Ssc_10019958 |
| Amazon molly | ENSPFOG00000010507 | Rockfish | Ssc_10021724 |
| Amazon molly | ENSPFOG00000010511 | | |
| Amazon molly | ENSPFOG00000010548 | | |
| Amazon molly | ENSPFOG00000010554 | | |
| Amazon molly | ENSPFOG00000010564 | | |
| Amazon molly | ENSPFOG00000010625 | | |
| Amazon molly | ENSPFOG00000024331 | | |

**Table S20.** Primers used in qPCR and ISH.

| Gene | Sequence (5' to 3') | Function |
|---|---|---|
| Ssc_10021724 | F: AATGTTGGAGCGGCTACTGC | qPCR |
| | R: GCCACGATGAGGAACAAAGC | qPCR |
| Ssc_10008384 | F: GATTCCCTTCACCATGAGCAGT | qPCR |
| | R: GACCTGTTTGCCTCCCGTT | qPCR |
| Ssc_10005765 | F: AGAACAACCAGCCCACCTTG | qPCR |
| | R: TGATGATGATGTTGATTCTCTTCCT | qPCR |
| Ssc_10001812 | F: GTCGTGGTGGTAAGCAGGTGG | qPCR |
| | R: CATGACTCTGATGTGGTTGTCCCT | qPCR |
| *Ssc-spag8* | F:ATTTAGGTGACACTATAGAAGAGAAATGGAGACTGTC ACTAC | ISH |
| | R:TAATACGACTCACTATAGGGAGATCAGTTGTCAAGTG GGA | ISH |

### Reference


1 S, K. *et al.* Canu: scalable and accurate long-read assembly via adaptive k-mer weighting and repeat separation. *Genome research* **27**, 722 (2017).

2 Walker, B. J. *et al.* Pilon: An Integrated Tool for Comprehensive Microbial Variant Detection and Genome Assembly Improvement. *Plos One* **9**, e112963 (2014).

3 Burton, J. N. *et al.* Chromosome-scale scaffolding of de novo genome assemblies based on chromatin interactions. *Nature Biotechnology* **31**, 1119 (2013).

4 Servant, N. *et al.* HiC-Pro: an optimized and flexible pipeline for Hi-C data processing. *Genome Biology* **16**, 259 (2015).

5 Langmead, B. & Pop, M. Ultrafast and memory-efficient alignment of short DNA sequences to the human genome. *Genome Biology* **10**, R25 (2009).

6 Durand, N. *et al.* Juicer Provides a One-Click System for Analyzing Loop-Resolution Hi-C Experiments. *Cell Systems* **3**, 95-98 (2016).

7 Harris, R. S. Improved Pairwise Alignmnet of Genomic DNA. (2007).

8 Zhao, X. & Hao, W. LTR_FINDER: an efficient tool for the prediction of full-length LTR retrotransposons. *Nucleic Acids Research* **35**, W265-W268 (2007).

9 J, J. *et al.* Repbase Update, a database of eukaryotic repetitive elements. *Cytogenetic and Genome Research* **110**, 462-467 (2005).

10 Benson, G. Tandem repeats finder: a program to analyze DNA sequences. *Nucleic acids research* **27**, 573-580 (1999).

11 Birney, E., Clamp, M. & Durbin, R. GeneWise and Genomewise. *Genome research* **14**, 988 (2004).

12 Grabherr, M. G. *et al.* Trinity: reconstructing a full-length transcriptome without a genome from RNA-Seq data. **29**, 644 (2011).

13 Kent, W. J. BLAT--the BLAST-like alignment tool. *Genome research* **12**, 656-664 (2002).

14 M, S. & S, W. Gene prediction with a hidden Markov model and a new intron submodel. *Bioinformatics* **19**, 215--225 (2003).

15 Elsik, C. G. *et al.* Creating a honey bee consensus gene set. **8**, R13 (2007).

16 Bairoch, A., . & Apweiler, R., . The SWISS-PROT protein sequence database and its supplement TrEMBL in 2000. *Nucleic Acids Research* **28**, 45 (2000).

17 Mulder, N. & Apweiler, R. InterPro and InterProScan: tools for protein sequence classification and comparison. *Methods in Molecular Biology* **396**, 59 (2007).

18 Kanehisa, M. & Goto, S. KEGG: kyoto encyclopedia of genes and genomes. *Nucleic acids research* **28**, 27-30 (2000).

19 Ashburner, M. *et al.* Gene Ontology: tool for the unification of biology. *Nature genetics* **25**, 25 (2000).

20 Li, H. & Durbin, R. *Fast and accurate short read alignment with Burrows–Wheeler transform*. (Oxford University Press, 2009).

21 Sim?O, F. A., Waterhouse, R. M., Panagiotis, I., Kriventseva, E. V. & Zdobnov, E. M. BUSCO: assessing genome assembly and annotation completeness with single-copy orthologs. *Bioinformatics* **31**, 3210-3212 (2015).

22 Langmead, B. & Salzberg, S. L. Fast gapped-read alignment with Bowtie 2. *Nature Methods* **9**, 357-359 (2012).

23 Patro, R., Duggal, G., Love, M. I., Irizarry, R. A. & Kingsford, C. J. N. m. Salmon provides fast and bias-aware quantification of transcript expression. **14**, 417 (2017).
24 Langfelder, P. & Horvath, S. J. B. b. WGCNA: an R package for weighted correlation network analysis. **9**, 559 (2008).
25 Oldham, M. *et al.* Functional organization of the transcriptome in human brain. *Nature Neuroscience* **11**, 1271 (2008).
26 Li, H. *et al.* TreeFam: a curated database of phylogenetic trees of animal gene families. *Nucleic Acids Research* **34**, D572 (2006).
27 Edgar, R. C. MUSCLE: multiple sequence alignment with high accuracy and high throughput. *Nucleic acids research* **32**, 1792-1797 (2004).
28 Stamatakis, A., ., Ludwig, T., . & Meier, H., . RAxML-III: a fast program for maximum likelihood-based inference of large phylogenetic trees. *Bioinformatics* **21**, 456-463 (2005).
29 Wu, T. *et al.* Bioinformatic analyses of zona pellucida genes in vertebrates and their expression in Nile tilapia. *Fish Physiology & Biochemistry* **44**, 1-15 (2018).
30 Lin, Q. *et al.* The seahorse genome and the evolution of its specialized morphology. *Nature* **540**, 395-399, doi:10.1038/nature20595 (2016).
31 Liu, X., Deng, Y., Ni, Y. & Li, Z. in *Design, Automation & Test in Europe Conference & Exhibition.*
